# N-acetyl-phenylalanine induces hepatic steatosis in MASLD by disrupting ER-mitochondria calcium coupling and mitochondrial lipid oxidation

**DOI:** 10.1101/2025.09.24.678188

**Authors:** Rémy Lefebvre, Théo Rousseaux, Nadia Bendridi, Stéphanie Chanon, Delphine Arquier, Alexandre Humbert, Nicolas Bertocchini, Bruno Pillot, Emmanuelle Meugnier, Aurélie Vieille-Marchiset, Margaux Nawrot, Claudie Pinteur, Sophie Ayciriex, Rohit Loomba, Jennifer Rieusset, Cyrielle Caussy

**Affiliations:** Univ Lyon, CarMen Laboratory, INSERM U1060, INRAE U1397, Université Claude Bernard Lyon 1, 69495 Pierre-Bénite and 69500 Bron, France; Lyon Hepatology Institute, IHU EVEREST, France; Institut des Sciences Analytiques, Univ Lyon, Université Claude Bernard Lyon 1, CNRS UMR 5280, Villeurbanne, France; MASLD Research Center, Division of Gastroenterology and Hepatology. University of California at San Diego, La Jolla, CA, USA; Hospices Civils de Lyon, Département Endocrinologie, Diabète et Nutrition, Hôpital Lyon Sud, 69495 Pierre-Bénite, France

**Author notes:** Co-first authors (Rémy Lefebvre, Théo Rousseaux). Co-last authors (Jennifer Rieusset, Cyrielle Caussy). **Corresponding authors:** <u>Pr. Cyrielle Caussy</u>, Service d’Endocrinologie, Diabète et Nutrition, Hôpital Lyon Sud, 165 Chemin du Grand Revoyet 69495, Pierre-Bénite CEDEX, France, <u>Dr. Jennifer Rieusset</u>, CarMeN Laboratory, IMR INSERM U1060/INRAE U1397, Hopital Lyon-Sud, secteur 2, Bâtiment CENS-ELI-2D, 165 chemin du Grand Revoyet, F-69310 Pierre Bénite, France.

**Keywords:** MASLD, hepatic steatosis, host-microbiota interplay, aromatic amino acids, N-acetyl-phenylalanine, mitochondria-associated membranes, calcium coupling, mitochondria, phenylacetic acid

## Abstract

**Background & Aims:** The gut-liver axis and hepatic ER-mitochondria miscommunication (at contact sites called MAMs) are involved in the development of metabolic dysfunction-associated steatotic liver disease (MASLD). We investigated the role of circulating aromatic amino acids (AAA) derived from phenylalanine and tyrosine in MASLD potentially through MAM alterations.

**Methods:** We analyzed AAA metabolomic profiles in individuals with and without MASLD and validated findings in a biopsy-proven cohort. The pro-steatogenic effect of MASLD-associated AAAs was validated *in vitro* using lipid labeling, MAM structural/functional assays, and palmitate-induced respiration. *In vivo* effects were tested in mice fed with candidate AAAs, and MAM involvement was confirmed by expressing a specific organelle linker *in vitro* and *in vivo*.

**Results:** N-acetyl-phenylalanine (NAPA) was strongly associated with hepatic steatosis and correlated with specific gut microbes. *In vitro*, NAPA promoted lipid accumulation by impairing ER-mitochondria calcium exchange via a LAT1-dependent electrogenic mechanism, reducing mitochondrial lipid oxidation. Chronic NAPA administration in mice induced steatosis and MAM disruption. Notably, enhancing ER-mitochondria contacts with an organelle linker prevented NAPA-induced steatosis *in vitro* and *in vivo*. Additionally, other phenylalanine- and tyrosine-derived AAAs reproduced NAPA’s effects, suggesting a class-dependent mechanism.

**Conclusion:** NAPA emerges as a MASLD-promoting metabolite, contributing to hepatic steatosis by disrupting ER-mitochondria calcium coupling and mitochondrial lipid oxidation.

**Lay Summary:** The gut-liver axis is a key component of the development of MASLD, and circulating gut-derived metabolites, notably AAAs derived from phenylalanine and tyrosine metabolism, have been associated with MASLD. However, the specific causal mechanisms of these AAA metabolites in MASLD development remain unexplored. Here, we identified NAPA, a gut microbiome linked metabolite elevated in MASLD patients, as a causal driver of hepatic steatosis both in vitro and in vivo. Mechanistically, NAPA alters ER-mitochondria calcium coupling leading to reduced mitochondrial lipid oxidation, highlighting a new mechanism with potential therapeutic implications.

**HIGHLIGHTS:**

- Circulating NAPA levels are increased in MASLD patients and correlate with hepatic steatosis.
- NAPA levels result from a complex host-microbiota interplay
- NAPA induces lipid accumulation by dampening ER-mitochondria calcium coupling and mitochondrial lipid oxidation.
- NAPA disrupts MAMs by a LAT1-mediated electrogenic mechanism.
- Other Phe- and Tyr-mediated metabolites have the same pro-steatogenic effect than NAPA pointing to a class-dependent effect.

## INTRODUCTION

The gut-liver axis is a key component of the development of metabolic dysfunction-associated steatotic liver disease (MASLD). Alterations in the gut microbiome have been observed across the full spectrum of MASLD including metabolic dysfunction-associated steatohepatitis (MASH), advanced fibrosis or cirrhosis [1–4]. These microbiome alterations may not solely result from this metabolic disorder, strongly linked to insulin resistance and obesity, but gut-derived metabolites appear to play a causal role in MASLD development [5, 6]. Several plasma microbial metabolites, have been linked to MASLD pathogenesis [7]. Notably, microbial products of aromatic amino acids (AAAs) from phenylalanine (Phe) and tyrosine (Tyr) metabolism have been associated with MASLD. Studies have identified metabolites like phenylacetic acid (PAA) and 3-(4-hydroxyphenyl)-lactate (HPL) or phenyl-lactate (PL) linked to hepatic steatosis and/or fibrosis [2, 8]. Plasma levels of Tyr and other AAAs correlate with liver fat content, further supporting the role of Phe and Tyr metabolism in MASLD [9]. However, the specific causal mechanisms of these microbial AAA metabolites in MASLD development remain unexplored.

Several studies also point to an involvement of mitochondria in the pathophysiology of MASLD. Morphological alterations, a reduction in oxidative and ATP synthesis capacities and an increase in the production of free radicals are observed during MASLD [10]. Interestingly, mitochondria are not isolated in cells, but they interact with the endoplasmic reticulum (ER) in contact sites, called mitochondria-associated membranes (MAMs), in order to regulate mitochondrial bioenergetics and cell signaling, in part via calcium coupling [11]. Consequently, MAMs play a crucial role in the regulation of metabolism and MAMs are considered as a hub of nutrients and hormone signaling, regulating hepatic metabolic flexibility and insulin sensitivity [12, 13]. Moreover, ER-mitochondria miscommunication was associated with hepatic insulin resistance and steatosis in different mice models and in humans [12, 14–17]. Importantly, we recently demonstrated that hepatic disruption of MAMs was an early event occurring during overnutrition and played a causal role in the development of hepatic insulin resistance and steatosis [14]. However, whether MAMs could be impacted by circulating Phe- and Tyr-derived AAA to influence hepatic steatosis during MASLD and by which mechanism is unknown.

By combining human clinical studies, *in vitro* experiments and preclinical approaches in mice, we aimed to explore the role of Phe and Tyr derived metabolites in the development of MASLD and the potential link with MAM alterations.

## MATERIALS AND METHODS

All material and methods have been described in details in the supplementary material.

### Human participants

We performed a cross sectional analysis of a prospective cohort study of 156 patients from the Twin and Family Study (ClinicalTrials.gov: NCT01643512) residing in Southern California, as previously described [8]. We then validated the findings from the derivation cohort by performing a cross-sectional analysis using the data of 156 participants with biopsy-proven NAFLD at the UCSD NAFLD Research Center [8].

### Animal models

Animal studies were performed in accordance with the French guidelines for the Care and Use of Laboratory Animals and approved by the regional ethic committee. Nutritional protocols and adenoviral infection are described in the supplementary material.

### Untargeted Metabolome profiling

Serum metabolite assessment was performed by *Metabolon, Inc* (Durham, NC, USA) in both the derivation and validation cohort.

### Gut-microbiome sequencing and analysis

The metagenomics analysis of the gut microbiome was performed using whole-genome shotgun sequencing of DNA extracted from stool samples in a subgroup of participants from the Twin and Family cohort with matching metabolomic assessment (n=69).

### NAPA Quantification

Circulating NAPA was quantified by targeted *LC-MS/MS Analysis*.

### Cell culture

In vitro experiments were performed in both Huh7 cells and primary mouse hepatocytes. For treatments, cells were cultured in basal (Bovine Serum Albumine, BSA) or lipotoxic (palmitate 100 µM) conditions, with metabolite (500 µM) supplementation or vehicle (0,5% ethanol) for controls.

### ER-mitochondria interactions

ER-mitochondria interactions were analysed either by *in situ* proximity ligation assay (PLA) or by transmission electronic microscopy (TEM).

### Lipid accumulation

Lipid accumulation was assessed either using BODIPY (493/503) lipid probe, oil red O (ORO) staining or by measuring triglycerides (TG) levels.

### Calcium imaging

All calcium measurements were performed, in the absence of extracellular calcium, after 36h of infection with Ad-4mtD3CPV overexpressing the FRET-based mitochondrial ratiometric calcium probe (Palmer et al., 2006).

### Mitochondrial respiration

Fatty acid oxidation (FAO)-related mitochondrial respiration was measured on intact PMH (500 000 hepatocytes) using the OROBOROS analyzer at 25°C.

### Gene expression analysis

Gene expression was quantified by reverse transcription followed by real-time PCR using a Rotor-Gene (QIAGEN).

### Statistical analysis

Data are expressed as the mean ± SEM. Statistical analyses were performed using GraphPad 10.1 and detailed in the figure legends.

## RESULTS

### Circulating NAPA is increased in MASLD patients and correlates with hepatic fat content and gut bacteria species

To identify relevant microbial AAA metabolites derived from Phe and Tyr metabolism associated with MASLD, we performed a secondary cross-sectional analysis of untargeted serum metabolomic assessment in a derivation cohort of 156 well-characterized participants with and without MASLD. Detailed baseline characteristics were previously published in Caussy et al.[8]; hepatic fat content was assessed using magnetic resonance imaging (MRI) proton density fat fraction (PDFF) and 36 participants had MASLD.

We identified several metabolites derived from Phe and Tyr metabolism with significantly higher levels in MASLD compared to non-MASLD and a few metabolites with lower levels in patients without MASLD (**Figure 1A**, **Table S1**). Among them, 12 metabolites were also significantly positively correlated with liver fat content measured by MRI-PDFF (**Figure 1B**, **Table S1**). To confirm the association between the identified AAA metabolites with hepatic steatosis, we next analyzed the correlation between histological grade of steatosis and AAA metabolites in an independent validation cohort of 156 unrelated participants with biopsy-proven MASLD cohort (**Figure 1B**, **Table S2**). The detailed baseline characteristics of the biopsy-proven MASLD has been previously described in Caussy et al.[8]. The distribution of histological grade of steatosis 0,1, 2 and 3 was 1.9%, 32.7%, 37.2% and 28.2%, respectively. Interestingly, only NAPA, PL and HPL were confirmed with positive correlation with hepatic steatosis. Overall, NAPA had the highest fold change in MASLD versus non-MASLD (**Figure 1C**) and the highest correlation with hepatic fat content (r:0.51,p<0.001,q<0.001, **Figure 1D**), with a confirmed significant increase in serum NAPA levels along with histological grade of steatosis (p=0.02, **Figure 1E**).

**Figure 1.**
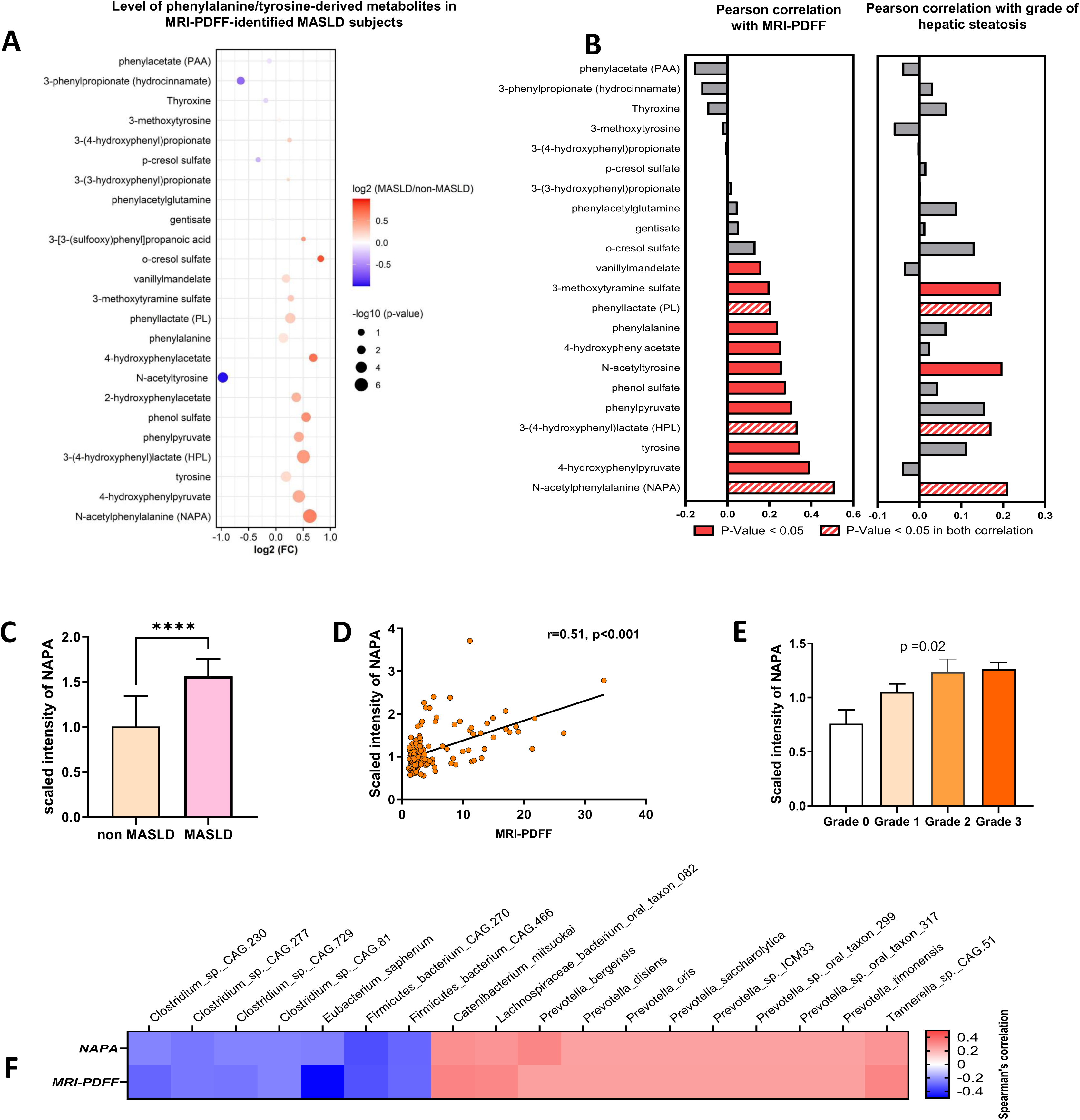
NAPA is increased in MASLD and associated with both hepatic fat content and gut-microbiome. (A) Bubble plot of serum level of Phe- and Tyr-derived metabolites as log2 fold-change in MASLD (n=36) versus non-MASLD (n=120), Welch’s t-tests. (B) Pearson correlation of these metabolites with hepatic fat (MRI-PDFF, Twin and Family cohort, left) or steatosis grade (biopsy-proven MASLD cohort, right). (C) Median scaled intensity ± 95%CI of NAPA in MASLD (n=36) and non-MASLD (n=120) (Welch’s t-test). (D) Pearson correlation of NAPA and MRI-PDFF in the Twin and Family cohort (n=156). (E) Median scaled intensity and 95% CI of NAPA level across histological grade of steatosis, in validation cohort with biopsy-proven MASLD (n=156) (Kruskal-Wallis test). (F) Heatmap of Spearman correlations between relative abundance of gut bacterial species NAPA or MRI-PDFF (Twin and Family subset n=69); adjusted for age and sex.

As NAPA is known to be a microbially produced metabolite [18], we thus performed a metagenomic analysis of the gut microbiome from the feces of human cohort. We found significant correlations between the relative abundance of bacterial species with both plasma NAPA concentration (**Table S3**) and hepatic fat content measured by MRI-PDFF (**Table S4**). Among them, 18 bacterial species were significantly positively or negatively correlated with both serum NAPA levels and MRI-PDFF, after adjusting for age and sex (**Figure 1F**), suggesting a complex interplay between host and microbiota.

### NAPA induces hepatic steatosis in vitro and in vivo in mice

To directly test the causal role of NAPA in the pathogenesis of MASLD, we treated Huh7 cells for 16 hours with NAPA (500µM), in both basal conditions or in co-treatment with palmitate (100µM), and measured the repercussions on cellular lipid accumulation using BODIPY staining. As shown in **Figure 2A**, NAPA treatment significantly induced lipid accumulation by increasing lipid droplets (LD) size (+18.5%,p<0.0001), without affecting LD number (Supplemental Figure S1A). Moreover, palmitate specifically increased LD size (+32.6%,p<0.0001) and co-treatment of cells with NAPA and palmitate induced a slightly higher lipid accumulation vs palmitate alone (+11%,p<0.0001) (**Figure 2A** and **Figure S1A**). We further confirmed the pro-steatotic effects of NAPA in more physiological primary mouse hepatocytes (PMH), as we also observed a significant lipid accumulation (+11,4%,p<0.0001), similarly to palmitate treatment (+6.7%,p<0.001), and a slightly additive effect when palmitate and NAPA are mixed vs palmitate alone (+6.2%,p<0.001) (**Figure 2B**). We also confirmed in PMH that the LD number was not modified by NAPA and/or palmitate treatments (**Figure S1B**).

**Figure 2.**
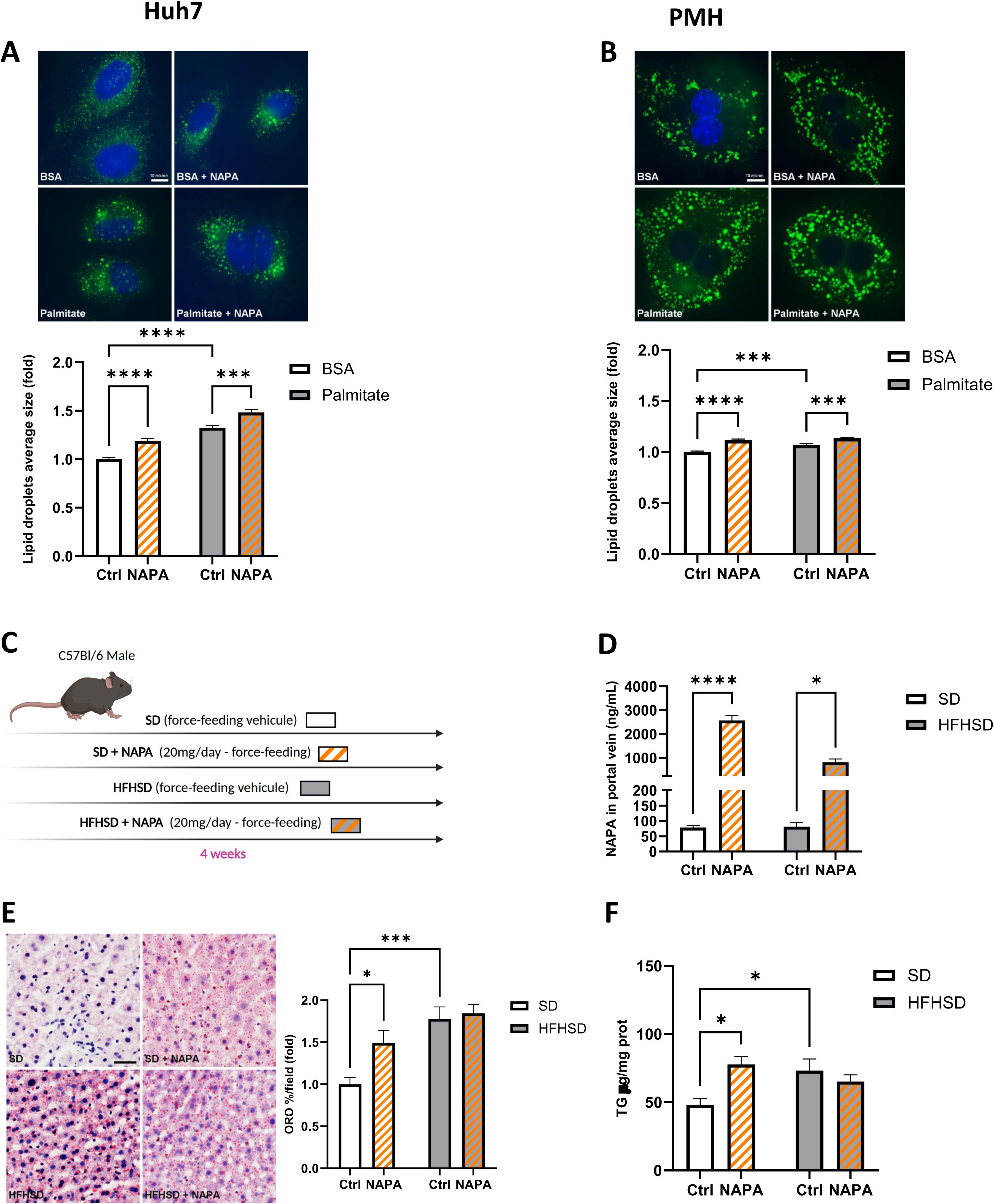
NAPA induces hepatic steatosis in vitro and in vivo in mice. (A-B) Huh7 cells (A) and PMH (B) treated for 16h with NAPA or vehicle in basal situation (BSA) or 100µM palmitate conditions. Representative images (top) and quantification (bottom) of lipid droplets average size by BODIPY in (A) Huh7 cells (N=3-6, n=113-126 cells) and (B) PMH (N=3, n=120 cells). (C) Schematic of *in vivo* experiment. Mouse phenotyping is shown in Figure S2. (D) Quantification of NAPA concentrations (LC-MS/MS,) in the portal vein of untreated and treated SD and HFHSD-fed mice (N=9 mice/group). (E) Quantification of hepatic lipid accumulation by ORO staining (N=9-12 mice/group, scale bar: 30μm). (F) Representative images and quantification of liver triglycerides (N=7-10 mice/group). Statistics: two Way ANOVA with Tukey’s multiple comparison test: NS: non-significant; *p<0.05; **p<0.01; ***p<0.001; ****p<0.0001.

To confirm these results *in vivo*, mice fed for four weeks with either a standard diet (SD_4w_) or a high-fat and high-sucrose diet (HFHSD_4w_) were force-fed mice with NAPA (20mg/mice) or vehicle (**Figure 2C**). NAPA treatment did not modify the body weight, the liver weight and the glucose tolerance of both SD_4w_ and HFHSD_4w_ (**Figure S2**). As expected, NAPA levels were significantly increased in the portal vein of NAPA-treated mice in both SD_4w_ and HFHSD_4w_, with a more pronounced increase in SD_4w_ (**Figure 2D**). Importantly, NAPA treatment induced hepatic lipid accumulation in SD_4w_ mice (+49.1%,p=0.038), as shown by the increase of ORO staining area of liver mouse section (**Figure 2E),** and confirmed by the increase of hepatic TG levels quantified from liver lysates (**Figure 2F**). As expected, HFHSD4w feeding also induced hepatic steatosis (+77.5%,p<0.001), whereas no additive effect of NAPA was observed in HFHSD_4w_ mice (**Figure 2E-F**). Altogether, these data show that NAPA is able *per se* to induce hepatic lipid accumulation both *in vitro* and *in vivo*.

### NAPA disrupts hepatic ER-mitochondria interactions in vitro and in vivo in mice

To further explore the involved mechanism, we investigated whether the pro-steatogenic NAPA effects are associated with a disruption of MAMs. For that, we used *in situ* PLA targeting the VDAC1-IP3R1 complex in fixed Huh7 cells, a reliable and thoroughly validated technique to quantify ER-mitochondria physical interactions [14, 16, 19]. As shown in **Figure 3A**, treatment of Huh7 cells for 16 hours with NAPA disrupted ER-mitochondria interactions, as illustrated by the reduction of VDAC1-IP3R1 dots per cell (-37.1%,p<0.0001). As previously observed [17], palmitate treatment also reduced ER-mitochondria interactions in 16h-treated Huh7 cells (-30.7%,p<0.0001), and co-treatment with metabolite and palmitate had few additive effects on MAMs vs palmitate alone (-24.63%,p<0.001) (**Figure 3A**). Importantly, we confirmed all these regulations in PMH, confirming the inhibitory effects of NAPA on ER-mitochondria interactions (**Figure 3B**). Lastly, we analyzed ER-mitochondria contact sites in the mouse liver of NAPA-treated SD_4w_ and HFHSD_4w_ mice, using both *in situ* PLA and transmission electronic microscopy (TEM) analyses. NAPA treatment significantly disrupted ER-mitochondria interactions in the liver of SD_4W_ mice (-42%,p=0.032) (**Figure 3C**). In agreement with previously published results [14], HFHSD_4w_ feeding also disrupted hepatic MAMs (-43%,p=0.02), and no additive effect of NAPA was observed in HFHSD_4w_ mice (**Figure 3C**). These data were confirmed by TEM analyses, as NAPA reduced the percentage of mitochondrial membrane in contact with the ER in SD_4W_ mice (-11.67%,p=0.021) (**Figure 3D**). A similar reduction was observed by HFHSD feeding (-11.46%,p=0.018), with no additive effect in presence of NAPA (**Figure 3D**). These results confirm that NAPA-induced hepatic lipid accumulation observed *in vitro* and *in vivo* is associated with disrupted ER-mitochondria interactions into hepatocytes.

**Figure 3.**
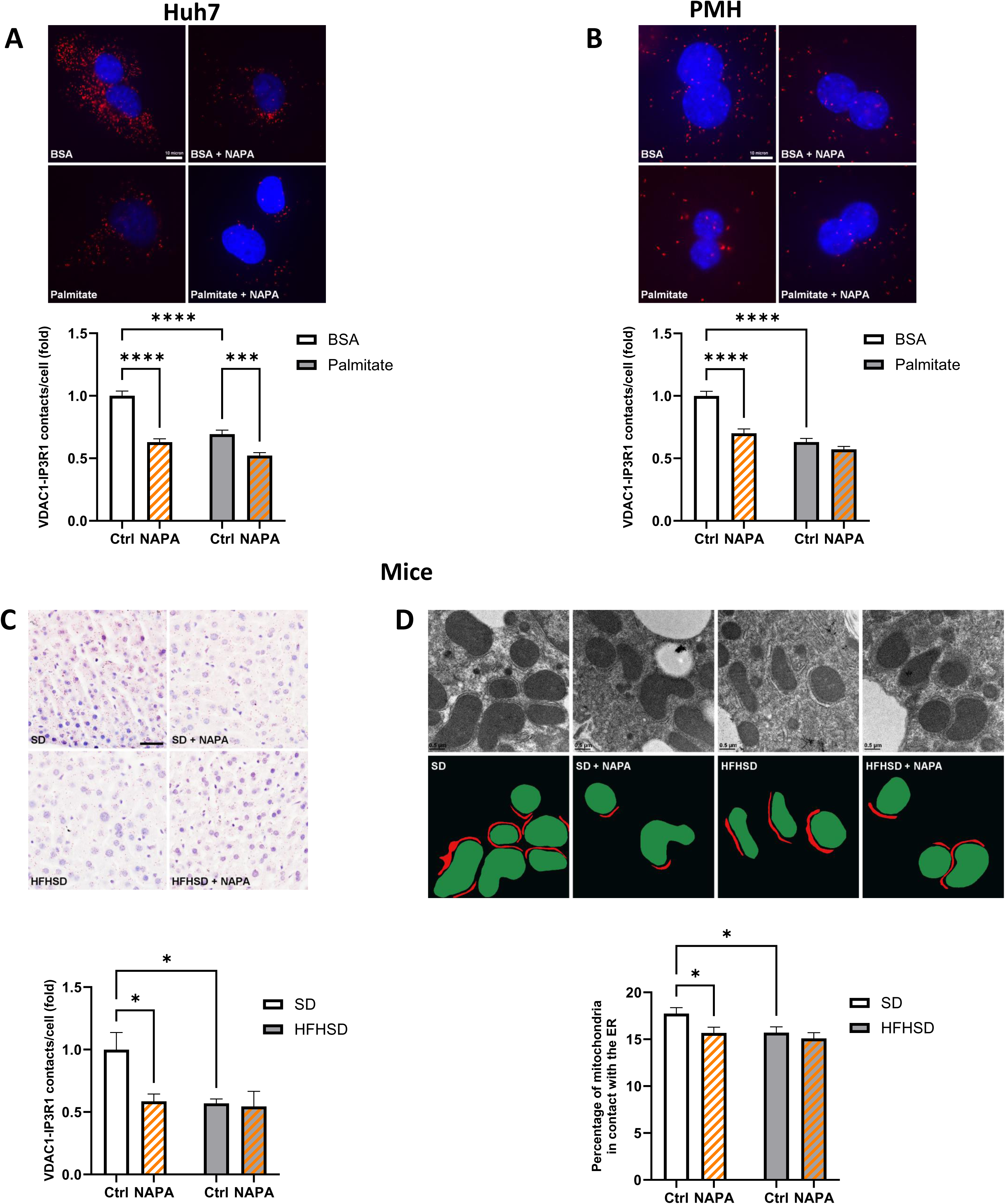
NAPA disrupts ER-mitochondria interactions both in vitro and in vivo. Huh7 cells (A) and PMH (B) were treated for 16h with NAPA or vehicle in basal situation (BSA) or 100µM palmitate condition. Representative images (top; scale bar, 10μm) and quantification (bottom) of IP3R1-VDAC1 dots/cell, measured by *in situ* PLA in (A) Huh7 cells (N=3, n=50-59 cells) and (B) PMH (N=3, n=90-101 cells). (C) Representative images (top, scale bar, 30μm) and quantification (bottom) of VDAC1-IP3R1 dots/cell measured, by *in situ* PLA on paraffin-embedded liver of untreated and NAPA-treated SD_4w_ and HFHSD_4w_ mice. (N=8-9 mice/group). (D) Representative images (top; scale bar, 0.5μm) and quantification of ER-mitochondria interactions using TEM analysis in mice experiments. The architecture of ER (red) and mitochondria (green) interactions were drawn graphically under TEM images; data were expressed as % MAMs/mitochondria in 50nm range (total interface) (N=6 mice/group and n= 263-334 mitochondria). Statistics: two way ANOVA (A-C) with Tukey’s multiple comparison test (D) or Fisher’s LSD test: NS, non-significant; *p<0.05; **p<0.01; ***p<0.001; ****p<0.0001.

### NAPA reduces ER-mitochondria calcium exchange and mitochondrial lipid oxidation in PMH

As ER-mitochondria interaction allows the exchange of calcium between both organelles, we imaged in NAPA-treated PMH the IP3R-mediated mitochondrial calcium accumulation using the adenovirus-expressed FRET-based 4mtD3CPV mitochondrial calcium sensor. As illustrated in **Figure 4A**, ATP stimulation increased mitochondrial calcium accumulation in 4mtD3CPV-expressing PMH, regardless of the treatment. However, NAPA treatment decreased basal mitochondrial calcium levels (-7.5%,p=0.0001, **Figure 4B**), and importantly also reduced ATP-stimulated mitochondrial calcium accumulation (-29%,p=0.0001, **Figure 4C),** indicating that NAPA disrupted ER-mitochondria calcium exchange.

**Figure 4.**
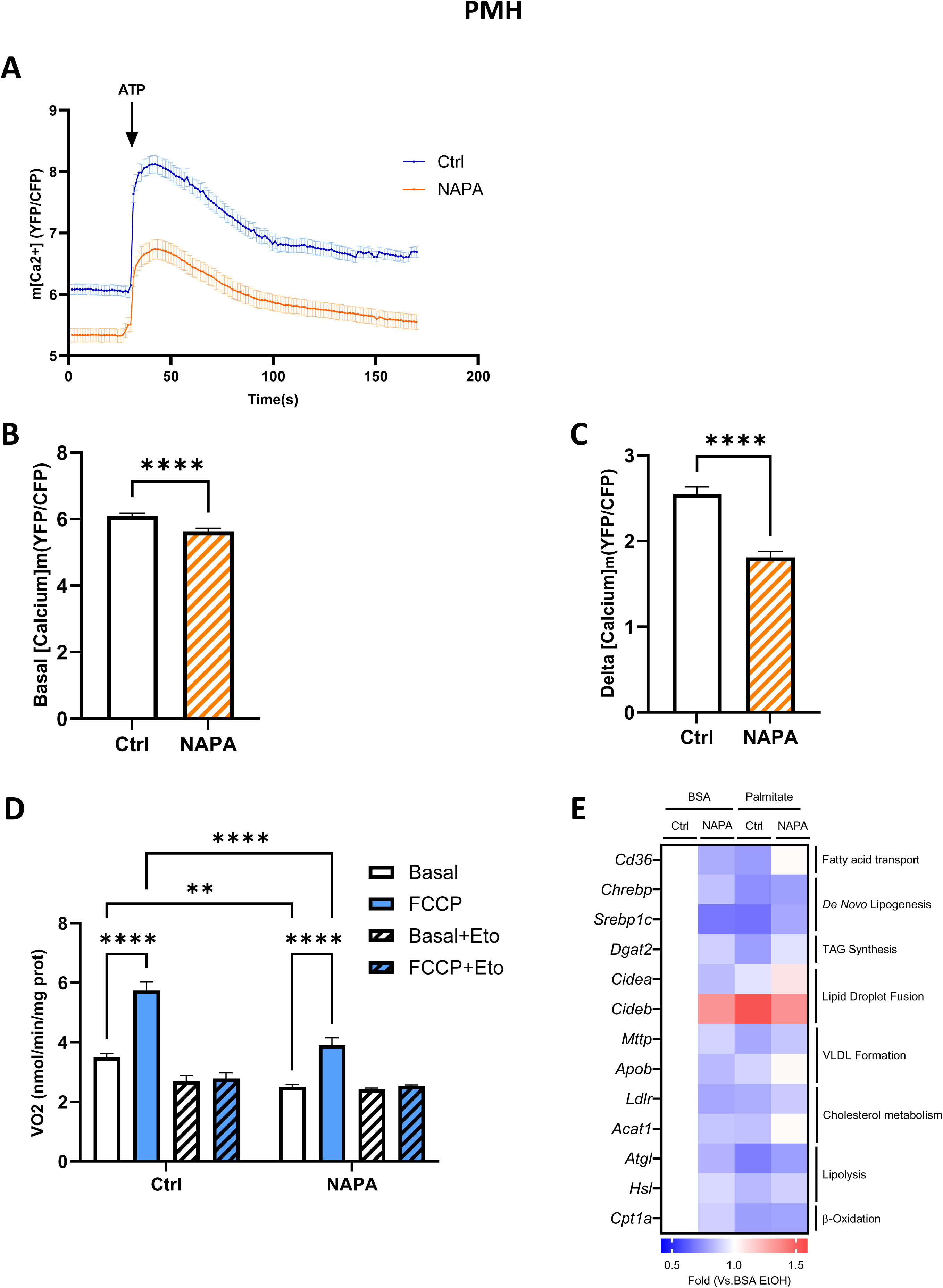
NAPA disrupts ER-mitochondria calcium exchange and reduces lipid-induced mitochondrial respiration. (A-C) Ad-4mtD3CPV-infected PMH were treated for 16h with NAPA or vehicle. (A) Illustration of calcium curves before and after ATP stimulation, (B-C) quantitative analysis of basal (B) and ATP-stimulated (C) mitochondrial calcium accumulation (N=8, n=111-124 cells, Mann-Whitney test). (D) PMH were treated for 16h with NAPA or vehicle. Basal and maximal FCCP-induced mitochondrial oxygen consumption were measured under palmitate in the absence or presence of etomoxir (Eto), an inhibitor of lipid oxidation (N=7, two-way ANOVA followed by Tukey’s multiple comparison test). (E) Heat map showing the regulation of genes involved in lipid metabolism in PMHs; data are expressed as fold change versus BSA-Ctrl condition (N=5, no significant statistical differences). Statistics: ANOVA followed with Tukey’s multiple comparison, ** p<0.01; ***p<0.001; ****p<0.0001.

Intra-mitochondrial calcium is crucial to control mitochondrial oxidative metabolism [20]. As hepatic lipid accumulation could be linked to reduced mitochondrial oxidation secondary to organelle miscommunication, we therefore measured using an OROBOROS system palmitate-linked mitochondrial oxygen consumption in intact PMH treated or not with NAPA. To that end, we assessed basal and maximal FCCP-induced mitochondrial oxygen using palmitate as substrate, in absence or presence of etomoxir, an inhibitor of lipid oxidation (n=8 independent PMH preparation). As shown in **Figure 4D**, NAPA treatment significantly reduced both basal respiration (-28.3%,p<0.01) and maximal FCCP-induced mitochondrial oxygen consumption (-31.9%,p<0.0001), whereas these effects are prevented by etomoxir treatment. NAPA may not impact other pathways of lipid metabolism, as mRNA levels of key genes of lipid import or export and of lipogenesis were not significantly impacted by NAPA (**Figure 4E**). Altogether, these data suggest that NAPA-induced lipid accumulation could be linked to reduced mitochondrial lipid oxidation, as a consequence of disrupted ER-mitochondria calcium coupling.

### Causal role of MAM disruption in NAPA-induced lipid accumulation

In order to investigate the causal role of MAM disruption on the pro-steatotic effect of NAPA, we first performed a kinetic study in Huh7 cells. After 1h of treatment of Huh7 cells with NAPA, we observed a significant reduction of ER-mitochondria interactions by *in situ* PLA (-33.1%,p<0.001, **Figure 5A**), suggesting an early effect of NAPA on MAMs, whereas the effect of NAPA on lipid accumulation was observed after at least 16 hours of treatment (**Figure 5B)**. These data indicate that the effect of NAPA on MAMs precedes the effect on lipid accumulation, in favor of a potential causal role of MAMs. We further investigated the causal role of MAMs by reinforcing experimentally ER-mitochondria interactions in Huh7 cells, using adenoviral-mediated overexpression of a synthetical organelle linker, as previously performed [14]. We confirmed by TEM analyzes that the linker increased ER-mitochondria interaction in untreated Huh7 cells and prevented the deleterious effects of NAPA on organelle contacts (**Figure 5C**). In addition, expression of the organelle linker prevented the effects of NAPA on lipid accumulation in Huh7 cells, whereas these effects were observed in Ad-Ctrl cells (**Figure 5D**).

**Figure 5.**
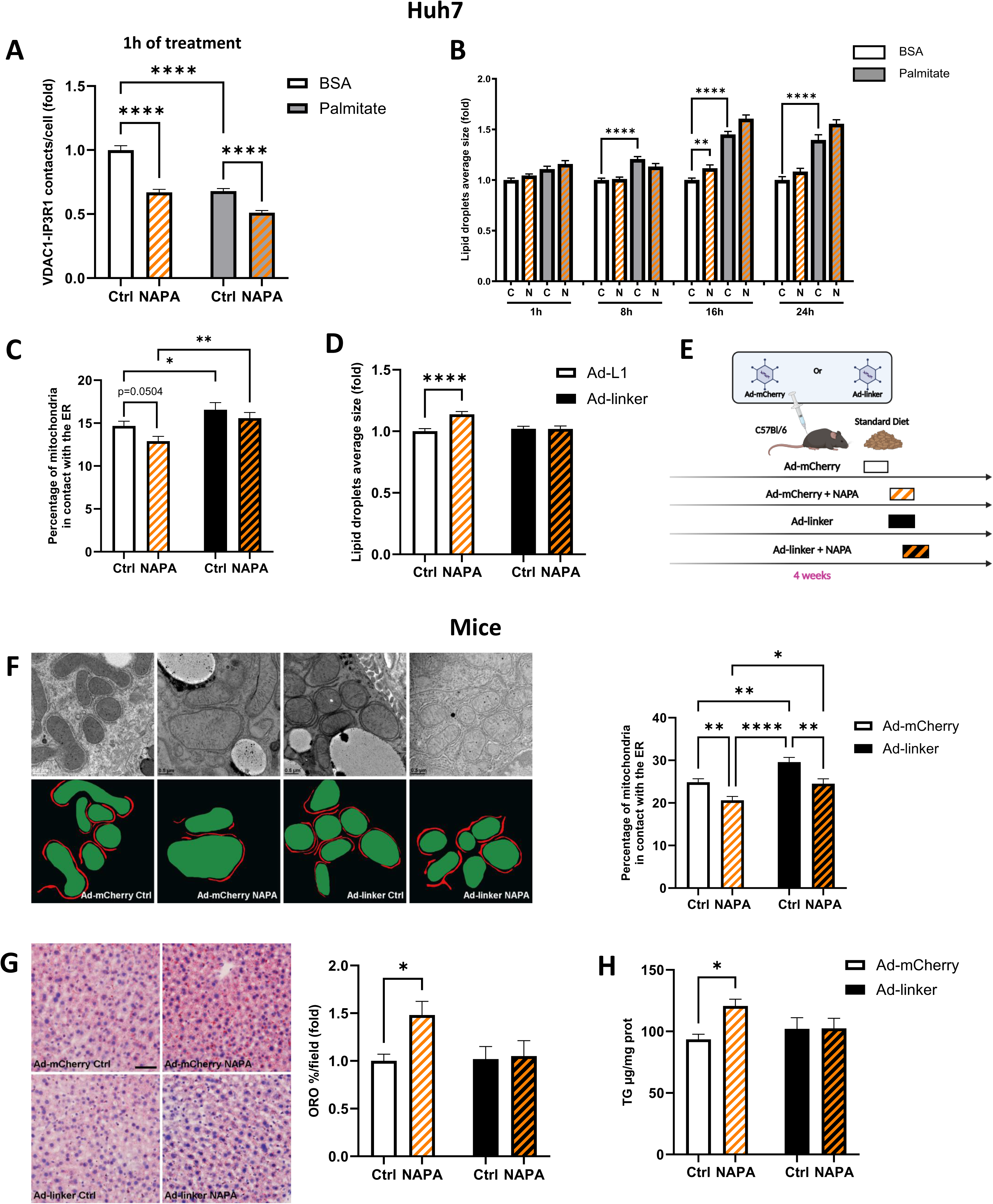
Causal role of disrupted MAMs in NAPA-induced lipid accumulation. (A-B) Kinetics experiments in Huh7 cells treated with NAPA or Ctrl under BSA condition or 100 µM palmitate conditions; (A) VDAC1-IP3R1 dots/cell, measured by *in situ* PLA, in Huh7 cells after 1h or treatment (N=3, n=83-90 cells). (B) Quantification of lipid droplets average size (BODIPY) after 1, 8, 16 or 24h of treatment (N=3, n=30 cells). (C-D) Huh7 cells were infected for 48h with adenovirus expressing either a control or the organelle linker. (C) ER-mitochondria interactions measured by TEM analysis; data are expressed as % MAMs/mitochondria in 50 nm range (total interface, N=3, n=145-167 mitochondria). (D) Lipid droplets average size measured by BODIPY (N=4, n=87-92 cells); (E) Schematic of in vivo experiment. (F) Representative TEM images (left, scale bar, 0.5μm) and quantification (right) of ER-mitochondria interactions in liver from infected mice. The architecture of ER (red) and mitochondria (green) interactions were drawn graphically under MET images. Data were expressed as % MAMs/mitochondria in 50 nm range (total interface, N=8 mice/group and n=185-257 mitochondria). (G) Representative images (left, scale bar, 46μm) and quantification (right) of hepatic lipid accumulation measured by ORO staining (N=7-8 mice/group). (H) Quantification of liver TG in infected mice (N=8 mice/group). Statistics: two-way ANOVA (A–D–F) with Tukey’s multiple comparison test, (B-G-H) or Fisher’s LSD test or (C) Mann-Whitney test. *p<0.05; **p<0.01; ***p<0.001; ****p<0.0001.

To confirm these data *in vivo,* we expressed the linker in the liver of SD4w mice, using adenovirus as previously performed [14], whereas control mice were infected with an Ad-mCherry adenovirus (**Figure 5E**). As expected, the expression of the linker increased ER-mitochondria interactions in the liver of mice, as illustrated by the increase of the total percentage of mitochondrial membrane in contact with ER measured by TEM (+19.2%,p<0.01, **Figure 5F**). Once again, NAPA treatment reduced ER-mitochondria interactions in Ad-mCherry mice (-17.08%,p<0.01); this effect remained significantly present in Ad-linker mice (-17.12%,p<0.01), however organelle interactions were higher compared to NAPA-treated Ad-mCherry mice (+18.97%,p<0.05) (**Figure 5F**). The presence of NAPA effect on MAMs in mice expressing the linker is likely mediated to a partial effect of the linker that may not have infected all the hepatocytes. Importantly, NAPA-induced hepatic steatosis was prevented by the expression of the linker, as illustrated by ORO staining (**Figure 5G**) and hepatic TG levels measurement (**Figure 5H**). Altogether, these data validate *in vitro* and *in vivo* that NAPA-induced hepatic lipid accumulation is dependent on the disruption of MAMs.

### NAPA effects on hepatic MAMs and lipid accumulation are mediated through a LAT1-mediated electrogenic mechanism

Next, we investigated the molecular mechanisms by which NAPA regulates MAMs. The cellular import of AAA through transporters, such as the L-type amino acid transporter 1 (LAT1), are known to close ATP-dependent K^+^ channel, inducing a membrane depolarization and the extracellular calcium entry through a voltage-dependent calcium channel [21, 22]. Thus, we investigated whether a similar mechanism contributes to NAPA effect on MAMs in hepatocytes, using complementary approaches (**Figure 6A**). Firstly, we inhibited LAT1 using either a specific siRNA reducing by 10 times the expression of *LAT1* (p<0.0005, **Figure S3A**) or the JPH203 inhibitor, a specific LAT1 inhibitor [23]. Genetic inhibition of LAT1 prevented the effects of NAPA on ER-mitochondria interactions (**Figure 6B**) and lipid accumulation (**Figure 6C**). These results were confirmed by treating Huh7 cells with JPH203 (**Figure S3B-C**). Secondly, we used either diazoxide to open, or tolbutamide to close the ATP-dependent K^+^ channel. Diazoxide treatment (100µM) of Huh7 cells for 16 hours prevented the disruptive effect of NAPA on MAMs (**Figure 6D**), whereas tolbutamide treatment (100µM) mimicked the effect of NAPA on MAMs by disrupting MAMs (**Figure 6F**). Then, diazoxide prevented the accumulation of lipids by NAPA (**Figure 6E**), whereas tolbutamide induced lipid accumulation (**Figure 6G**). Thirdly, we used EGTA (10mM) or BAPTA (10 µM), to chelate extracellular or intracellular calcium respectively. We found that 1 hour of treatment of Huh7 cells with EGTA or BAPTA prevented the effects of NAPA on MAMs (**Figure 6H-I**). We did not measure the repercussions on lipid accumulation, because calcium chelators were toxic during longer incubation. Collectively, these data indicate that NAPA disrupts ER-mitochondria interaction, at least in part, through an increase of cytoplasmic calcium following a LAT1-mediated electrogenic effect.

**Figure 6.**
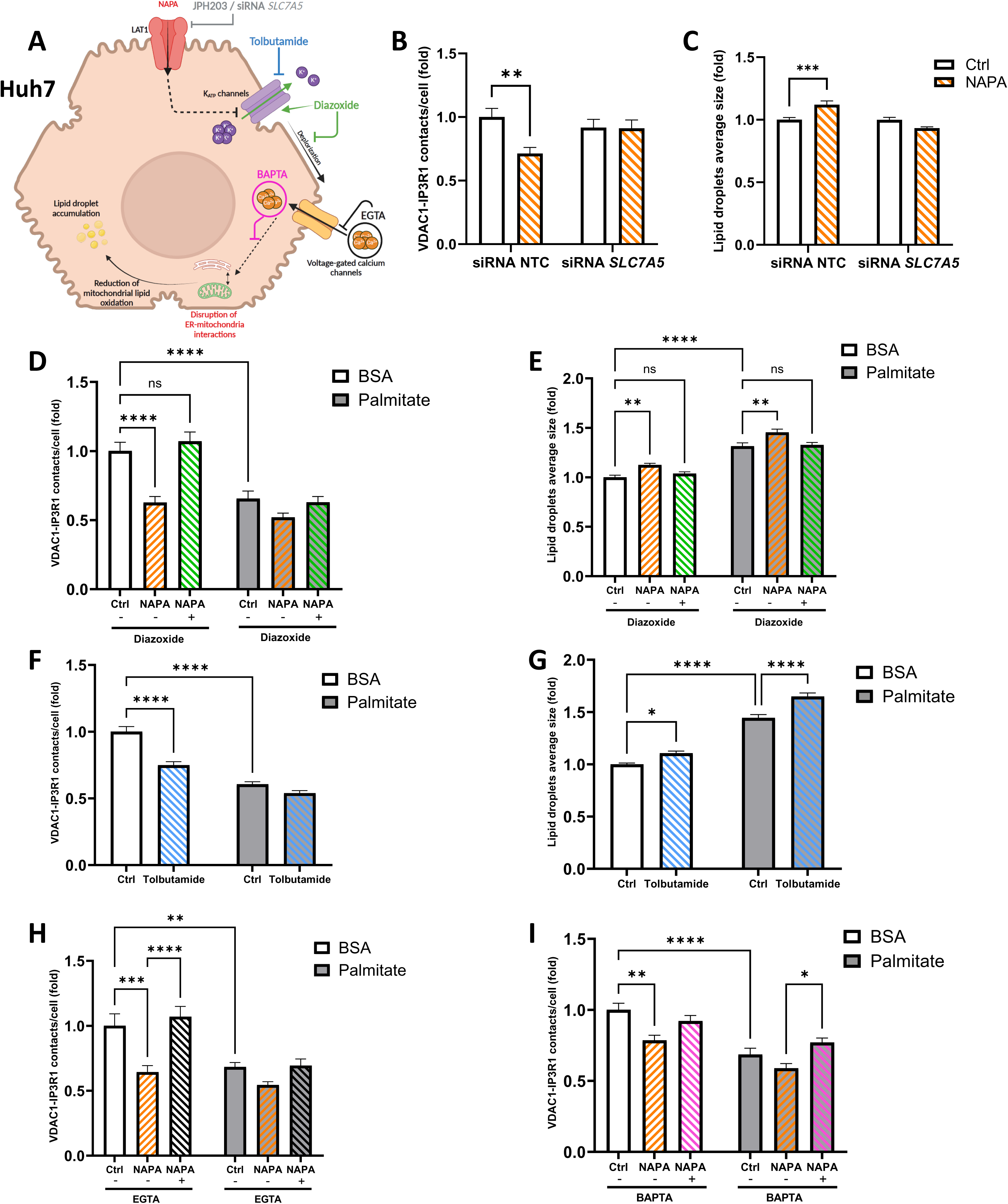
NAPA disrupts MAMs and induces lipid accumulation through a LAT1 electrogenic effect. (A) Graphical representation of NAPA transporter and key pathway regulators. (B-C) Huh7 cells transfected with siRNA-NTC or siRNA-SLC7A5 for 24h, then treated 16h with NAPA or Ctrl. (B) quantification of IP3R1-VDAC1 dots/cell by *in situ* PLA; (C) lipid droplets average size (BODIPY). (N=2 in triplicate, n=59-66 cells). (D-I) Huh7 cells treated with NAPA or Ctrl ± diazoxide, tolbutamide, EGTA, or BAPTA, with or without palmitate (100 µM, 16h (D-G) or 1h (H-I); (D-F, H-I) IP3R1–VDAC1 PLA quantification (N=1–3, n=20–30 cells), (E, G) lipid droplet size (BODIPY) (N=1–2, n=40–67 cells). Statistics: two-way ANOVA with Tukey’s multiple comparison test: NS, non-significant; *p<0.05; **p<0.01; ***p<0.001; ****p<0.0001.

### Other Phe- and Tyr-derived AAAs disrupt organelle communication and induce lipid accumulation

Our clinical assessment also identified HPL and PL as elevated in MASLD and strongly correlated with hepatic fat content in both cohorts. Like NAPA, these AAA-derived metabolites also disrupted MAMs in both Huh7 cells (**Figure S4A and S4E**) and PMH (**Figure S4B and S4F**). In addition, HPL and PL also induced lipid accumulation in both Huh7 cells (**Figure S4C and S4G**) and PMH (**Figure S4D and S4H),** without any significant change in the number of lipid droplets **(Figure S4)**.

As PAA was previously associated with hepatic steatosis in both humans and mice by unknown mechanism [2], we investigated whether PAA could induce lipid accumulation in a MAM-dependent pathway. Consistent with NAPA effects, PAA treatment significantly disrupted ER-mitochondria interaction and induced lipid accumulation in both Huh7 cells and PMH (**Figure 7A - 7**). In PMH, PAA reduced basal and ATP-stimulated mitochondrial calcium accumulation (**Figure 7E-F**), confirming that PAA disrupts ER-mitochondria calcium coupling. In addition, PAA treatment significantly reduced both basal and maximal palmitate-related mitochondrial respiration, whereas these effects were prevented by etomoxir treatment (**Figure 7G**), suggesting that PAA, like NAPA reduces mitochondrial lipid oxidation. Moreover, the effect of PAA was MAM-dependent, as the expression of the linker in Huh7 cells reduced the inhibitory effect of PAA on MAMs (**Figure 7H**) and importantly prevented PAA-induced lipid accumulation (**Figure 7I)**. Lastly, mice fed for 4 weeks SD and HFHSD with admixed 0,8% PAA (**Figure 7J**), showed reduced ER-mitochondria interactions in the liver (**Figure 7K**), and increased hepatic lipid accumulation (**Figure 7L-M**). Collectively, these data suggest that the effects of Phe- and Tyr-derived metabolites on hepatic MAMs and steatosis is class-dependent.

**Figure 7.**
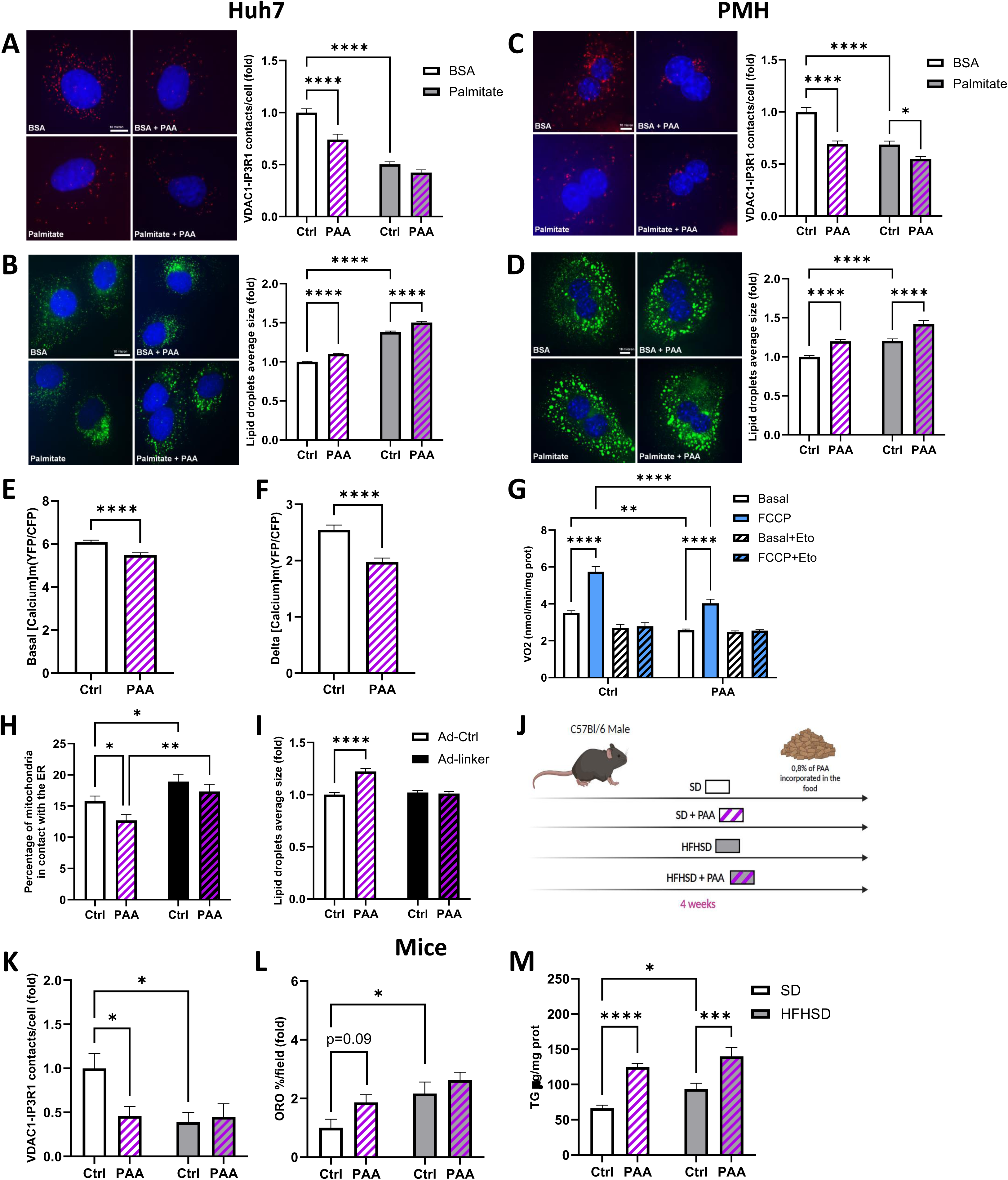
Effect of PAA in lipid accumulation and integrity of MAMs. (A-I) Huh7 cells (A-B, E-I) or PMH (A, C) were treated for 16h with PAA or vehicle under standard condition (BSA) or 100 µM palmitate. (A, C) Representative images (left) and quantification (right) of IP3R1-VDAC1 dots/cell using *in situ* PLA in Huh7 (A, N=3, n=83-87 cells) and PMHs (C, N=3-4, n=60cells). (B, D) Quantification of lipid droplet average size by BODIPY in Huh7 cells (B, N=3-6, n= 210 cells) and PMH (D, N=3-4, n=79-80 cells). (E-F) Quantification of basal (E) and ATP-stimulated (F) mitochondrial calcium accumulation (N=8, n=98-111 cells). (G) Basal (white) and maximal (blue) FCCP-induced mitochondrial oxygen consumption measured in PMHs under palmitate ± etomoxir (Eto), an inhibitor of lipid oxidation (N=7). (H) ER-mitochondria interactions measured by TEM analysis in infected Huh7 cells; data are expressed as % MAMs/mitochondria in 50 nm range (total interface, N=3, n=74-96 mitochondria). (I) Quantification of lipid droplets average size in infected Huh7 (BODIPY), (N=3, n=87 cells). (J) Schematic of in vivo experiment (N=8-9 mice/group). (K) Quantification of VDAC1-IP3R1 dots/cell, by *in situ* PLA, on paraffin-embedded mouse liver. (L) Hepatic lipid accumulation by ORO staining. (M) Liver triglyceride content Statistics: two-way ANOVA (A-D, G, I, K) with Tukey’s multiple comparison test, (H, L-M) or Fisher’s LSD test, or (E-F) by Mann-Whitney test: *p<0.05; **p<0.01; ***p<0.001; ****p<0.0001.

## DISCUSSION

By combining untargeted serum metabolomics data from two independent cohorts and mechanistic experiments *in vitro* and *in vivo*, we demonstrated here the causal role of some Phe- and Tyr-derived AAAs in the pathogenesis of MASLD. Indeed, NAPA, HPL and PL were strongly associated with hepatic steatosis in both human cohorts and all three metabolites induced lipid accumulation in both Huh7 cells and PMH. Here, we highlighted the high relevance of NAPA as a novel biomarker for MASLD at an early stage before the development of fibrosis. Interestingly, PAA, another Phe-derived AAA with pro-steatogenic effects, was not significantly associated with MASLD in our cohorts but was previously associated with hepatic steatosis in non-diabetic women with obesity [2], pointing potential differences in environmental factors, geography, or gut-microbiome composition. In agreement with this study [2], we confirmed that PAA induces lipid accumulation in Huh7 cells and PMH, as well as *in vivo* in 4 week-treated mice. However, we went one-step further by demonstrating that the pro-steatogenic effect of PAA is associated with the alteration of MAMs, both *in vitro* and *in vivo*.

The origin of circulating Phe- and Tyr-derived metabolites is only partially understood. While they could be produced by mammalian metabolism [24], NAPA, PAA, HPL and PL may also have a microbial origin and may be derived from AAA metabolism catalyzed by the gut-microbiome [2, 8]. We have previously linked the levels of HPL and PL to the abundance of bacterial species significantly associated with MASLD [3]. Similarly, PAA was associated with changes in the gut-microbiome associated with MASLD [2]. Here, we identified several bacterial species in the human gut-microbiome that were significantly correlated positively or negatively with NAPA levels and liver fat content, suggesting either production or consumption of NAPA by the bacterial species. In agreement, Dekkers et al. [25] reported several significant positive and negative correlations between plasma levels of NAPA and bacterial species from the human gut-microbiome, including a positive correlation between *Catenibacterium mitsuokai* and NAPA, also observed in our study (Figure 1F). Interestingly, the metabolomic profile of culture media from a large panel of individual bacterial strains available online (https://sonnenburglab.github.io/Metabolomics_Data_Explorer) reported that *Catenibacterium mitsuokai* produced NAPA [18]. Similarly, species of the family *Prevotellaceae*, significantly associated with plasma NAPA concentration and intrahepatic fat content in our study, produced NAPA. Conversely, other species from the *Clostridiacea* family induced a reduction in NAPA in the culture media suggesting a consumption of the NAPA or its precursor. This could also explain the negative correlation found in this study between some bacterial species from *Clostridiacea* Family and NAPA and intrahepatic fat content. This highlights the complexity of the microbiome-host interactions for the regulation of circulating NAPA. Further investigations, beyond the scope of this study, are now required to better elucidate NAPA metabolic pathways in the context of MASLD.

The mechanisms by which Phe- and Tyr-mediated metabolites induce hepatic steatosis were until now completely unknown. We showed that NAPA, PAA, HPL and PL disrupt hepatic ER-mitochondria interactions measured by *in situ* PLA in Huh7 cells and PMH. For NAPA and PAA, we confirmed their effect on MAMs by TEM showing that they disrupt ER-mitochondrial calcium coupling and reduce mitochondrial lipid oxidation in PMH. Furthermore, we confirmed *in vivo* that 4 weeks of treatment with NAPA or PAA is sufficient to disrupt ER-mitochondria interactions in the mouse liver (confirmed by both *in situ* PLA and MET), accompanied by the induction of hepatic steatosis. Although we only tested the effect of NAPA and PAA *in vivo*, the fact that HPL and PL consistently disrupted MAMs *in vitro* as well, suggests that similar results should be observed *in vivo*, with possible additive effects. Finally, we validate the causal role of MAM disruption in hepatic steatosis, as increasing MAMs in the liver of mice prevents NAPA-induced hepatic steatosis. Interestingly, we also confirmed this causal role of disrupted MAMs in hepatic steatosis with PAA-induced hepatic steatosis *in vitro*. Overall, we demonstrated that hepatic MAMs can sense circulating Phe- and Tyr-derived metabolites in portal blood and adjust ER-mitochondrial calcium exchange to control mitochondrial oxidative metabolism.

Regarding the mechanisms by which AAA metabolites affect MAMs, we demonstrated here the involvement of a LAT1-mediated electrogenic mechanism and an increase in intracellular calcium levels leading to a disruption of MAMs, at least for NAPA and PAA. Further studies are now required to determine how elevation of intracellular calcium levels disrupts ER-mitochondria interactions in hepatocytes. AAA have been reported to act on specific receptors/transporters or ‘sensor’ mechanisms that are then able to initiate cell signaling events and regulate intracellular events. These AAA transporters often act by co-transporting some ions (Na+, K+ or H+), which is consistent with the electrogenic effect we observed [26]. For NAPA, its electrogenic effect may be mediated by the essential AA transporter LAT1 which is a neutral amino acid transporter that is sodium independent, since genetic and pharmacological inhibition of LAT1 prevents the effect of NAPA on MAMs. Interestingly, LAT1 has been suggested to act as an ‘environmental sensor’ of amino acid availability and may be involved when AAA levels increase due to gut-microbiome dysbiosis [26]. As LAT1 is not very specific, this could potentially explain why all AAA tested in our study had the same effect on MAMs and hepatic steatosis.

In conclusion, our study provides 3 original conceptual advances: i) we identify NAPA as a novel pro-MASLD metabolite whose serum elevation in MASLD patients is related to a complex host-microbiota interplay, ii) we demonstrate that NAPA induces hepatic steatosis by disrupting ER-mitochondria calcium coupling and reducing mitochondrial lipid oxidation, and iii) we show that ER-mitochondria communication could be downregulated by AAA metabolite by a LAT1-mediated electrogenic mechanism.

## FINANCIAL SUPPORT AND ACKNOWLEDGMENT

This work was supported by INSERM and the national research agency (ANR-21-CE14-0088-01 from JR). RéL was supported by a research fellowship from the French government of higher education and research. RoL receives funding support from NCATS (5UL1TR001442), NIDDK (U01DK061734, U01DK130190, R01DK106419, R01DK121378, R01DK124318, P30DK120515), NHLBI (P01HL147835), and John C Martin Foundation (RP124). We thank the CIQLE platform from Claude Bernard University (Lyon) for TEM analyses, Gyorgi Hajnoczky and Gyorgi Csordas (Thomas Jefferson institute, Philadelphia, USA) for the generous gift of the ER-mitochondria linker and for sharing their macro to analyse ER-mitochondria interactions by TEM, Lexane Brunet (lab assistant) for her help with MET analysis, Guillaume Zoulim (PhD student) for his help for drawing Figures 1A and 1B, and Sandra Pesenti (lab assistant for her help in genomics analysis). Théo Rousseaux was supported by a grant from IHU EVEREST ANR-23-IAHU-0008. Rémy Lefebvre was supported by a research fellowship from the French government of higher education and research.

## AUTHORS CONTRIBUTION

RéL and TR conducted *in vitro* experiments and helped with the *in vivo* protocol, performed analysis and interpretation of the data, and drafted the manuscript. NaB conducted mice experiments, SC and AVM performed histological analysis on the mouse liver. DA and SA performed NAPA assessment in mice models. AH, MN and CP helped with some in vivo experiments. NiB performed calcium experiments and BP helped to isolate primary mouse hepatocytes. EM performed gut-microbiome statistical analysis. RoL: designed and funded the clinical cohort, recruited and phenotyped patients, collected biological samples, interpretation of data, performed critical revision of the manuscript. JR and CC performed study concept and design, analyzed and interpreted data, drafted of the manuscript, critical revision of the manuscript. JR and CC are the guarantors of this work and, as such, had full access to all the data in the study and took responsibility for the integrity of the data and the accuracy of the data analysis.

All authors approved the final version of this article.

---

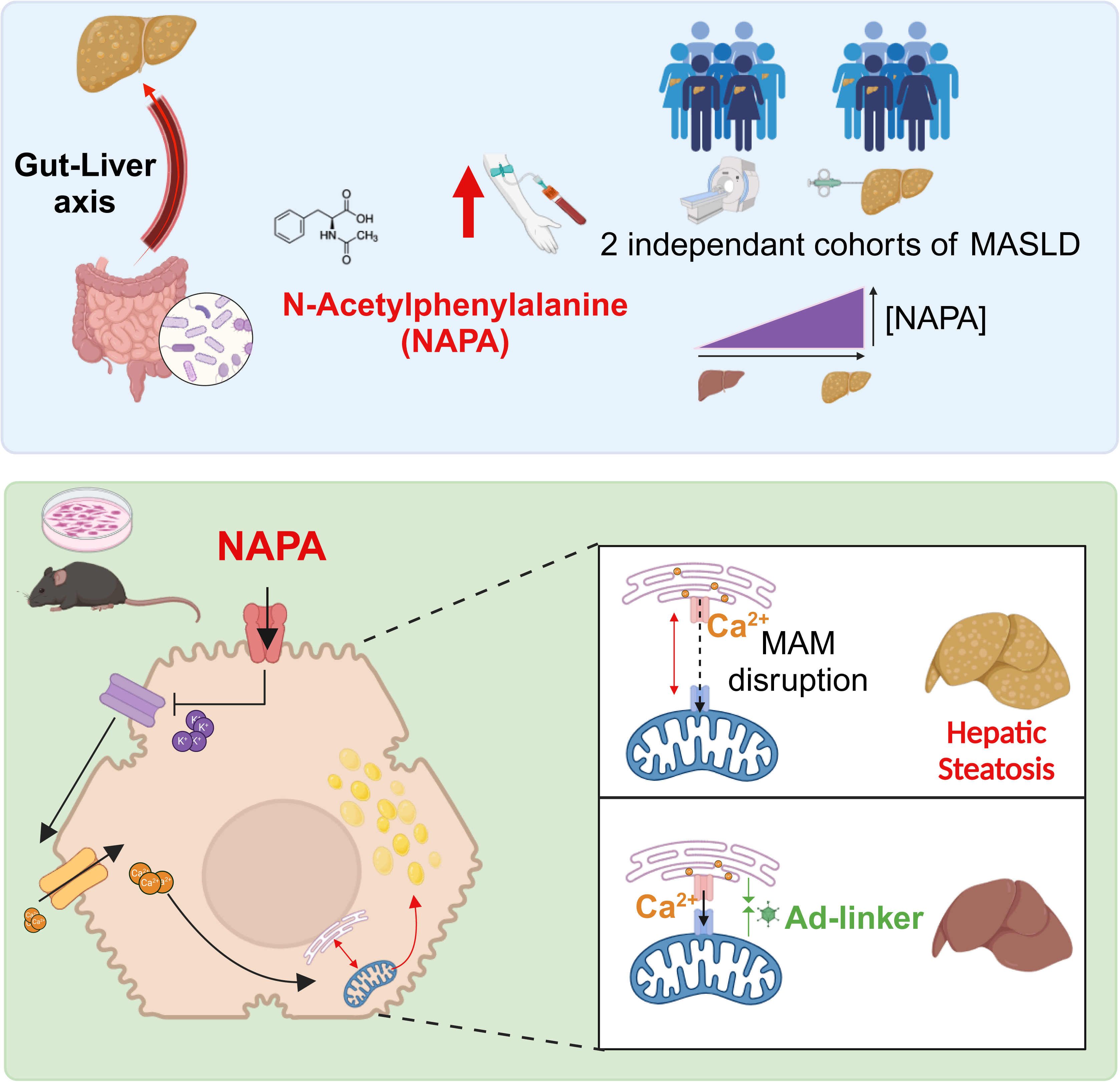

