## Supplemental methods and figures for "N-acetyl-phenylalanine induces hepatic steatosis in MASLD by disrupting ER-mitochondria calcium coupling and mitochondrial lipid oxidation"

**EXPERIMENTAL MODEL AND STUDY PARTICIPANT DETAILS**

### Human participants

Study participants and design of the derivation cohort form the Twin and Family Study.

We performed a cross sectional analysis of a prospective cohort study of patients from the Twin and Family Study (ClinicalTrials.gov: NCT01643512) residing in Southern California. This study included a total of 156 participants, 100 twins (50 twin pairs) and 56 participants either siblings or parents-offspring recruited by the University of California at San Diego (UCSD) NAFLD Research Center [1] between December 2011 and January 2014. Research visits and imaging procedures were performed the same day for each pair of twins, parent-offspring or siblings. This study was Health Insurance Portability and Accountability Act (HIPAA) compliant and was approved by the UCSD Institutional Review Board approval number: 111282. Informed written consent was obtained from each participant before enrolling in the study. Detailed inclusion and exclusion criteria have been previously described in Caussy et al.[1]. The presence of MASLD was defined by MRI-PDFF and participants with MRI-PDFF < 5% were considered as non-MASLD. Detailed MRI protocol has been previously described in Caussy et al. [1].

Study participants of the validation cohort of patients with biopsy-proven MASLD.

We have validated the findings from the derivation cohort by performing a cross-sectional analysis using the data of 156 participants with biopsy-proven NAFLD prospectively recruited between October 2011 and May 2014 at the UCSD NAFLD Research Center [1]. All patients with suspected NAFLD with a clinical indication for liver biopsy underwent a careful evaluation for other causes of hepatic steatosis and liver disease. This study was Health Insurance Portability and Accountability Act (HIPAA) compliant and was approved by the UCSD Institutional Review Board. Informed written consent was obtained from each participant before enrolling in the study. Detailed inclusion and exclusion criteria have been previously described in Caussy et al. [1].

***Animal models***

C57Bl/6J-Ola Hsd mice (Envigo, France): Animal studies were performed in accordance with the French guidelines for the Care and Use of Laboratory Animals and approved by the regional ethic committee (CECCAP-LS-2021-003). The mice were maintained at 25 °C and on an artificial 12-h light–dark cycle, with the light period beginning at 7:30 am. 5-week-old male mice were housed in groups of 6 on sawdust bedding in plastic cages in an enriched environment. Mice were fed with either a standard diet (SD, 35.6% carbohydrates, 10.2% fat by mass, Genobios, France) or a high-fat, high-sucrose diet (HFHSD, 260-HF from SAFE, 33.65% carbohydrate, 35.85% fat, 22.8% protein, in mass) for 4 weeks. In addition, mice were force-fed daily with 20 mg N-acetyl-L-phenylalanine (NAPA) or vehicle for 4 weeks. For PAA treatment, 0,8% of PAA was incorporated in the food (SAFE) and given to mice during 4 weeks. For the organelle linker protocol, 4-week-old male mice were infected (retro-orbital injection, $10^9$ infection-forming units/mouse) with recombinant adenovirus expressing either Ad-mCherry (as control) or Ad-Linker, and were forced-fed daily with 20 mg NAPA or vehicle for 4 weeks. At the end of each protocol, overnight- or 6h-fasted mice were weighed, anaesthetized with ketamine/xylasine (2/1) before blood was drawn from the portal vein, and the liver was collected, fixed or frozen in liquid nitrogen.

### METHOD DETAILS

#### *Untargeted Metabolome profiling*

Serum metabolite assessment was performed by *Metabolon, Inc* (Durham, NC, USA) in both the derivation and validation cohort. Samples were extracted and split into equal parts for analysis on the GC/MS and LC/MS/MS platforms [2]. Software was used to match ions to an in-house library of standards for metabolite identification and for metabolite quantitation by peak area integration [3]. For further details regarding serum metabolites data acquisition have been previously provided in Caussy et al. [1]. Missing values, if any, were imputed with the observed minimum for that particular compound.  Statistical comparisons including Welch's t-tests to compare between the MASLD and non-MASLD groups and analyses for MRI-PDFF measurements and histological grade of steatosis were carried out, along with Pearson correlations to assess relationships between clinical parameters and metabolite levels. Statistical analyses were performed on natural log-transformed data. Data are presented in scaled intensity where each biochemical in the original scale was re-scaled to have median equal to 1. Statistical analyses were performed using ArrayStudio and the programs R. The False Discovery Rate using the Benjamini-Hochberg method was used to account for multiple testing.

***Gut-microbiome sequencing and analysis***

The metagenomics analysis of the gut microbiome was performed using whole-genome shotgun sequencing of DNA extracted from stool samples in a subgroup of participants from the Twin and Family cohort with matching metabolomic assessment (n=69). The detailed protocol for the DNA extraction, library preparation and sequencing has been described previously in Loomba et al. [4]. The partial correlation between the relative abundance of gut-microbiome species and logNAPA and with hepatic fat content measured by MRI-PDFF controlling for age and sex were performed using ppcor package (v1_1) with R (v4.2.1)

***NAPA Quantification in blood from mice models***

*Sample Preparation:* 50 µL of blood was mixed with 50 µL of cold $H_2O$/MeOH (1:4, *v/v*) and vortexed for 30 sec. Extracts were diluted with 200 µL of cold $H_2O$. Samples were stored for 5 min at -20°C and were centrifuged for 15 min at 13300 rpm. 100 µL of the supernatant were transferred into a vial and then analyzed by targeted LC-MS/MS.

*Targeted LC-MS/MS Analysis*: The chromatographic separation of NAPA was carried out on a Xselect CSH C18 column (3,5 µm, 2.1 × 100 mm, Waters, Milford, USA). The mobile phase A was composed of $H_2O$/CAN (85:15, *v/v*) with 10 mM ammonium acetate and 0.1% formic acid and the mobile phase B was composed of IPA/ACN/$H_2$0 (90:9,5:0,5, *v/v/v*) with 10 mM ammonium acetate and 0.1% formic acid. The initial conditions of the gradient were 1% of phase B. The phase B content was increased up to 99% during 24,26 min and kept for 2 min. Next, phase B content was decreased to 1% for re-equilibration from 27.26 min up to 30 min. The total run time of the method was 30 min. The flow rate was 0.5 mL/min, the injection volume was 2 µL, and the column temperature was 70 °C. The NAPA analysis was performed in positive ion mode with the following parameters: the ion source gas 1 was 50 psi, the ion source gas 2 was 50 psi, the curtain gas was 35 psi, the CAD gas was set to 9 unit, the capillary temperature was set to 550°C, spray voltage was 5500V. A pause time was set to 5 msec. The LC–MS/MS of NAPA was operated in the MRM mode under unit mass resolution in both the Q1 and Q3 mass analyzers. The dwell time was set to 150 msec. Data acquisition and processing was performed using Sciex OS software (v.3.0).

***Cell culture***

Huh7 cells: cells were cultured in DMEM 1 g/L glucose supplemented with 10% FBS and 1% antibiotics, at 37°C and 5% CO2. For treatments, cells were cultured for 1h to 24h in basal (Bovine Serum Albumine, BSA) or lipotoxic (palmitate 100 µM) conditions, with metabolite (500 µM) supplementation or vehicle (0,5% ethanol) for controls. For organelle linker experiments, cells were infected with recombinant adenovirus expressing Ad-Ctrl (mAKAP1-mRFP, as a control) or Ad-Linker (mAKAP1-mRFP-yUBC6) for 48h prior treatments. For electrogenic experiments, cells were co-treated with diazoxide (100 µM), EGTA (10 mM), BAPTA (10 µM) or treated with tolbutamide (100 µM). Cells were also co-treated with LAT1 inhibitor Nanvuranlant/JPH203 (1 µM).

Primary murine hepatocytes (PMH): PMH were isolated using collagenase perfusion through the portal vein. The percentage of intact cells was determined by the trypan blue exclusion method. PMH were plated on collagen type I-coated culture plates and cultured in DMEM with 3 g/L glucose at 37°C and in a humid atmosphere with 5% $CO_2$. After 4h, the medium was replaced with fresh one during 24h before treatments. DMEM was supplemented with 2 mM L–glutamine, 1% penicillin streptomycin solution, 7.5 mM DL-sodium lactate, and 10% fetal bovine serum (FBS). After 24h hours, cells were cultured for 16h in basal (BSA) or lipotoxic (palmitate 100 µM) condition with metabolite (500 µM) supplementation or vehicle (ethanol) for controls.

***In situ* Proximity Ligation Assay (PLA)**

ER-mitochondrial interactions were assessed using *in situ* PLA targeting reticular 1,4,5-triphosphate receptor (IP3R1) (071213M, Merck Millipore France) and the mitochondrial voltage-dependent anion channel 1 (VDAC1) (ab14734, Abcam, UK). For cells, the assays were conducted on 4% paraformaldehyde-fixed and 0.1% triton-permeabilized Huh7 cells and PMH using a red-fluorescent in situ PLA DUOLINK kit (Merck, Darmstadt, Germany). For tissues, the assays were carried on in 4 µm paraffin sections of 10% formaldehyde-fixed and paraffin-embedded mouse liver samples, using a bright-field in situ PLA DUOLINK kit (Merck, USA). Fluorescence was analyzed with a Zeiss inverted fluorescent microscope using the AxioVision program. Quantification of signals was done with the BlobFinder software (Centre for Image Analysis, Uppsala University) and expressed as the number of blobs per nucleus compared to control.

***Transmission Electron Microscopy***

For ultrastructural study, cells and tissues were fixed with 2% glutaraldehyde (EMS) in 0.1 M sodium cacodylate (pH 7.4) buffer at 4°C. After washing three times in 0.2 M sodium cacodylate buffer, they were post-fixed with 1% aqueous osmium tetroxide (EMS) at room temperature for 1 hour and contrasted with tea Oolong in cacodylate 0.2 M pH 7.4 (EM grade) for 1 hour at room temperature. After washing three times in 0.2M sodium cacodylate and 1 time in water, they were dehydrated in a graded series of ethanol at room temperature and embedded in Epon. After polymerization, ultrathin sections (100 nm) were cut on a UC7 (Leica) ultramicrotome and collected on Cupper grids 150 mesh. Sections were stained with lead citrate before observations on a Jeol 1400JEM (Tokyo,Japan) operating at 100kV transmission electron microscope equipped with an Orius 600 camera (2688x2672 pixels,32-bits grayscale images) and Digital Micrograph (product version 1.7). For each sample, 11 to 20 pictures at 25,000x magnification were taken. ER and mitochondria were delimited using the Fiji® software and the fraction of mitochondrial membrane in contact with ER within a 50nm range was calculated and normalized to the mitochondrial perimeter and expressed as total % of organelle contact. Results are the mean of all analyzed mitochondria. The architecture of interactions between ER and mitochondria in the TEM images were drawn graphically using GIMP software 2.10.38.

***Lipid accumulation***

Lipid accumulation in Huh7 and PMH was assessed using BODIPY (493/503) lipid probe. Fluorescence was analyzed with a Zeiss inverted fluorescent microscope using the AxioVision program and lipid droplets average size and number were quantified using a custom written Fiji macro. For the mouse liver, lipid accumulation was evaluated using Oil Red O staining (ORO-k-250, Biognost) in 10µm-cryosections. In addition, hepatic lipid accumulation in the mouse liver was also analyzed by measuring triglycerides (TG) levels on liver lysates by fluorimetry (Triglyceride-Glo™ Assay, Promega).

***Calcium imaging***

All calcium measurements were performed, in the absence of extracellular calcium, in the following buffer (NaCl 1.4 mM; KCl 50 mM; MgCl2 10mM; HEPES 100mM; glucose 100mM EGTA 1mM, pH 7.4), after 36h of infection with Ad-4mtD3CPV overexpressing the FRET-based mitochondrial ratiometric calcium probe [5]. After 2-min basal fluorescence measurement, Na-ATP (100 µM, added in puff) was added to stimulate ER-mitochondria calcium exchange and acquisitions were continued during 2 min on 1 field/dish. The fluorescence ratio YFP/CFP was analyzed with MetaFluor 6.3 (Universal Imaging) after removing background fluorescence. Results represent the average of all analyzed cells in 3 dishes/condition from 8 independent experiments.

***Mitochondrial respiration***

Mitochondrial oxygen consumption rate was measured on intact PMH (500 000 hepatocytes) using the OROBOROS analyzer at 25°C. For fatty acid oxidation (FAO)-related respiration, intact PMH were incubated in FAO assay buffer (111mM NaCl, 4.7mM KCl, 1.25mM CaCl2, 2 mM MgSO4, 1.2 mM NaH2PO4), supplemented with 2.5 mM glucose, 0.5 mM carnitine and 5mM Hepes (pH 7.4). Kinetic studies were performed by sequential addition of palmitate (200µM), oligomycin (2 µM), FCCP (25 µM), and rotenone/antimycine A (0.5 µM/4 µM). Etomoxir (40 µM) was used to inhibit FAO. Results are expressed as both basal and FCCP-stimulated maximal respiration measured in 8 independent PMH preparation.

***Gene expression analysis***

Total RNA from Huh7 cells was extracted by using the TRI Reagent Solution (SIGMA). Levels of specific mRNA were quantified by reverse transcription followed by real-time PCR using a Rotor-Gene (QIAGEN). A standard curve was systematically generated with different amounts of purified target cDNA, and each assay was performed in duplicate and normalized using TATA-binding protein (TBP) mRNA level, as previously reported. Primers are listed in Table S5.

**QUANTIFICATION AND STATISTICAL ANALYSIS**

Data are expressed as the mean ± SEM. Statistical analyses were performed using GraphPad 10.1. Comparisons between more than 2 groups were analyzed using ANOVA followed by Tukey's, or Fisher's LSD multiple comparison test. *In vitro* experiments were performed in 1 to 6 independent experiments in triplicate (=N). *In vivo* experiments were performed in N=9-12 mice/group. For single cell imaging analysis, statistics were performed on n=number of cells to assess single cell effect as well as heterogeneity between them. Both N and n values, as well as the statistical test, are indicated in the figure legends. Significance was defined as a value of $p<0.05$.

### SUPPLEMENTAL FIGURES

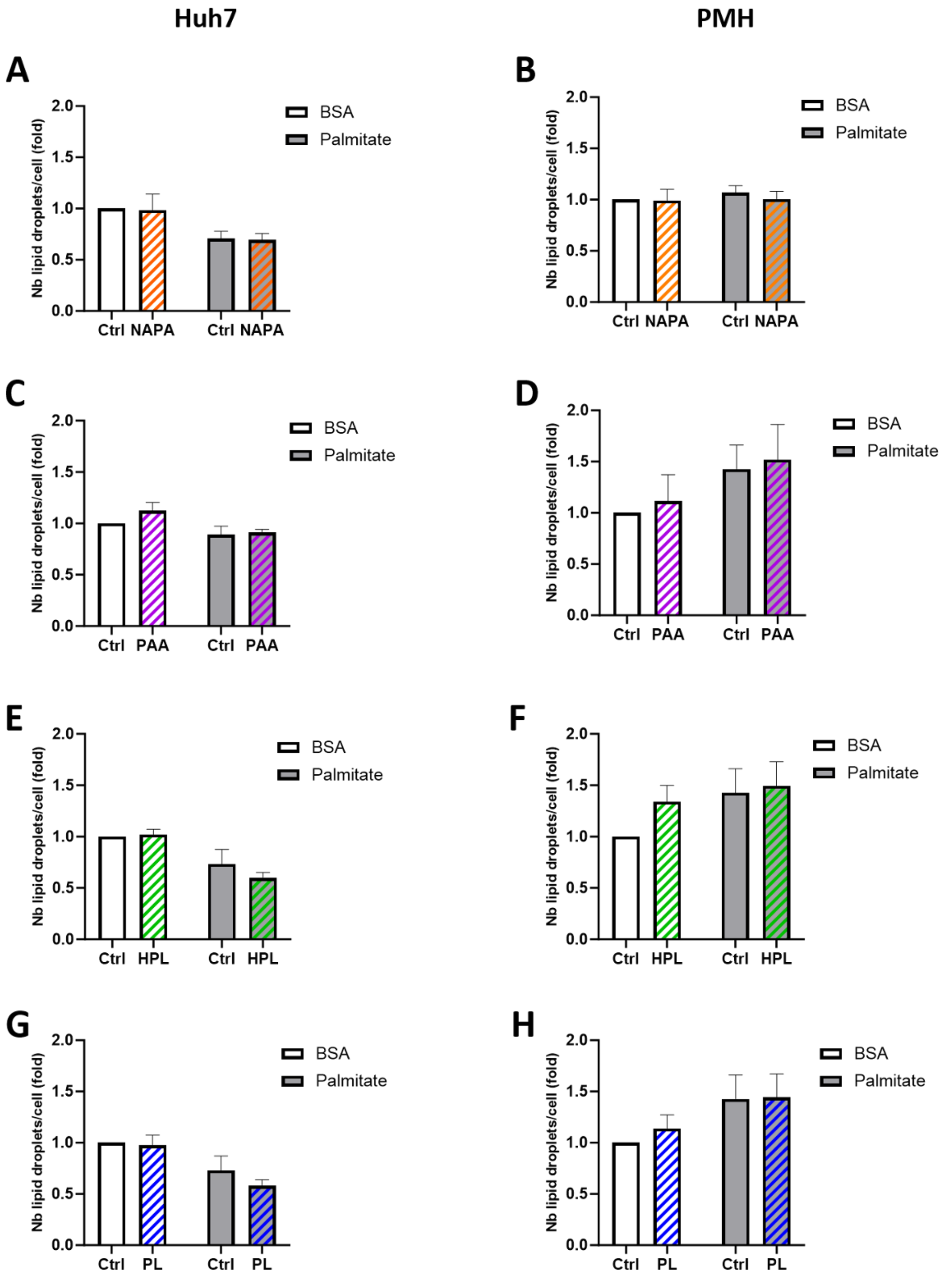


**Figure S1: Effects of Phe- and Tyr-derived metabolites on lipid accumulation are independent of variation in lipid droplets number.**

(A-H) Huh7 cells (A, C, E, G) or primary mouse hepatocytes (PMH), (B, D, F, H) are treated with metabolites for 16h. Quantitative analysis of lipid droplets number in both Huh7 cells (A, C, E, G) (N=3-6 experiments, n=91-176 cells) and PMH (B, D, F, H) (N=3 experiments, n=66 cells) measured by BODIPY staining. Results are expressed as a fold of control and number per cell. Statistical analysis: Two Way Anova followed by Tukey’s multiple comparison test: no comparison is significant.

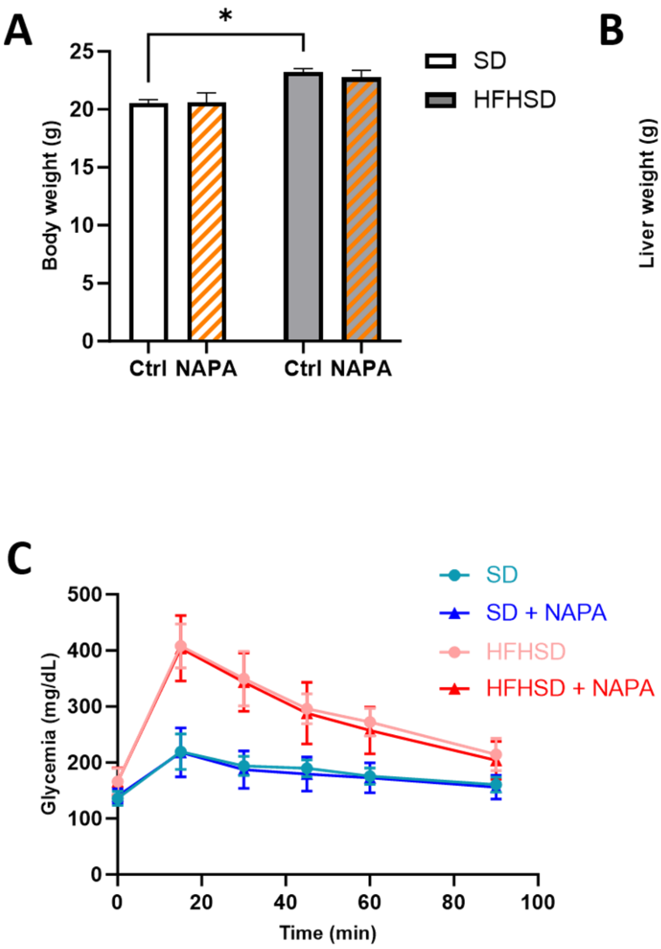


**Figure S2: Phenotyping data of SD and HFHSD mice treated with NAPA for 4 weeks.**

(A-C) Male mice were put on either a standard diet (SD, white bar) or a high-fat and high-sucrose diet (HFHSD, grey bar) for 4 weeks and force-fed daily with NAPA (orange dashed bar) or vehicle (empty bar). Graphical representation of the mouse experiment was illustrated in Figure 2C. Repercussion of treatments on body weight (A), liver weight (B) and glucose tolerance test (C) of mice. Statistical analysis: Two Way Anova followed by Tukey's multiple comparison test: *p <0.05.

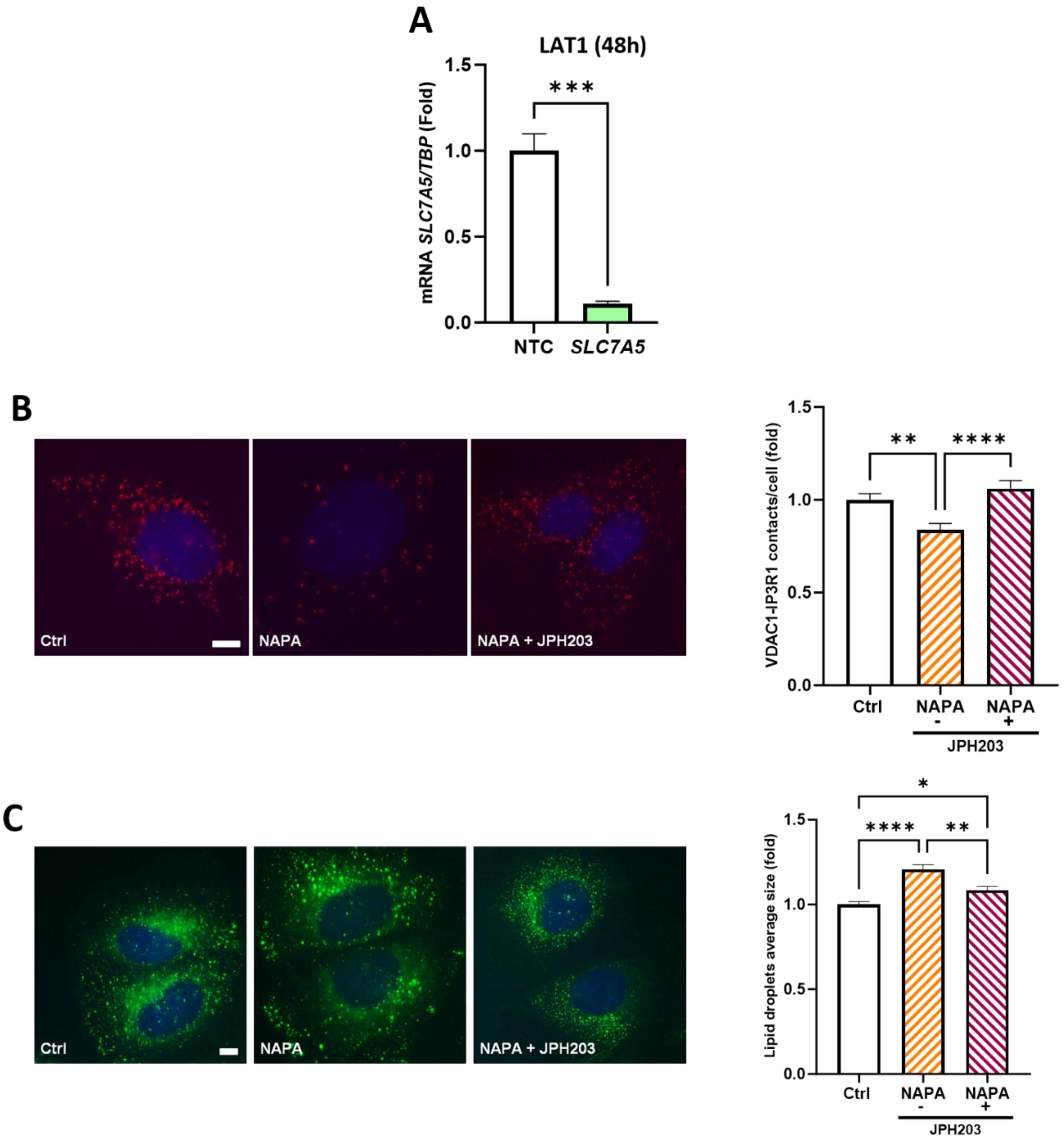


**Figure S3: LAT1 inhibition by JPH203 prevents NAPA effects on MAMs and steatosis in Huh7 cells.**
(A) Effects of the SLC7A5 siRNA on the mRNA levels of LAT1 transporters in Huh7 cells after 48h of silencing. Values are normalized by TBP and reported to the control (N=3 samples/condition; n=2 replicates per sample). (B-C) Huh7 cells were treated for 16h with NAPA (dashed bar) or vehicle (white bar), in absence (orange) or presence (red) of the LAT1 inhibitor JPH203. (B) Representative images (left, scale bar: 10 µm) and quantitative analysis (right) of IP3R1-VDAC1 dots/cell, measured by *in situ* PLA (N=3 experiments, n=77-80 cells). Results are expressed as a fold of control. (C) Representative images (at left) and quantitative analysis (at right) of lipid droplet average size measured by BODIPY staining (N=3 experiments, n=60 cells). Results are expressed as a fold of control. Statistical analysis: (B) Unpaired T-Test: ***p<0.001; (C-D) One Way Anova followed by Tukey’s multiple comparison test: *p <0.05; **p<0.01; ***p <0.001; ****p <0.0001.

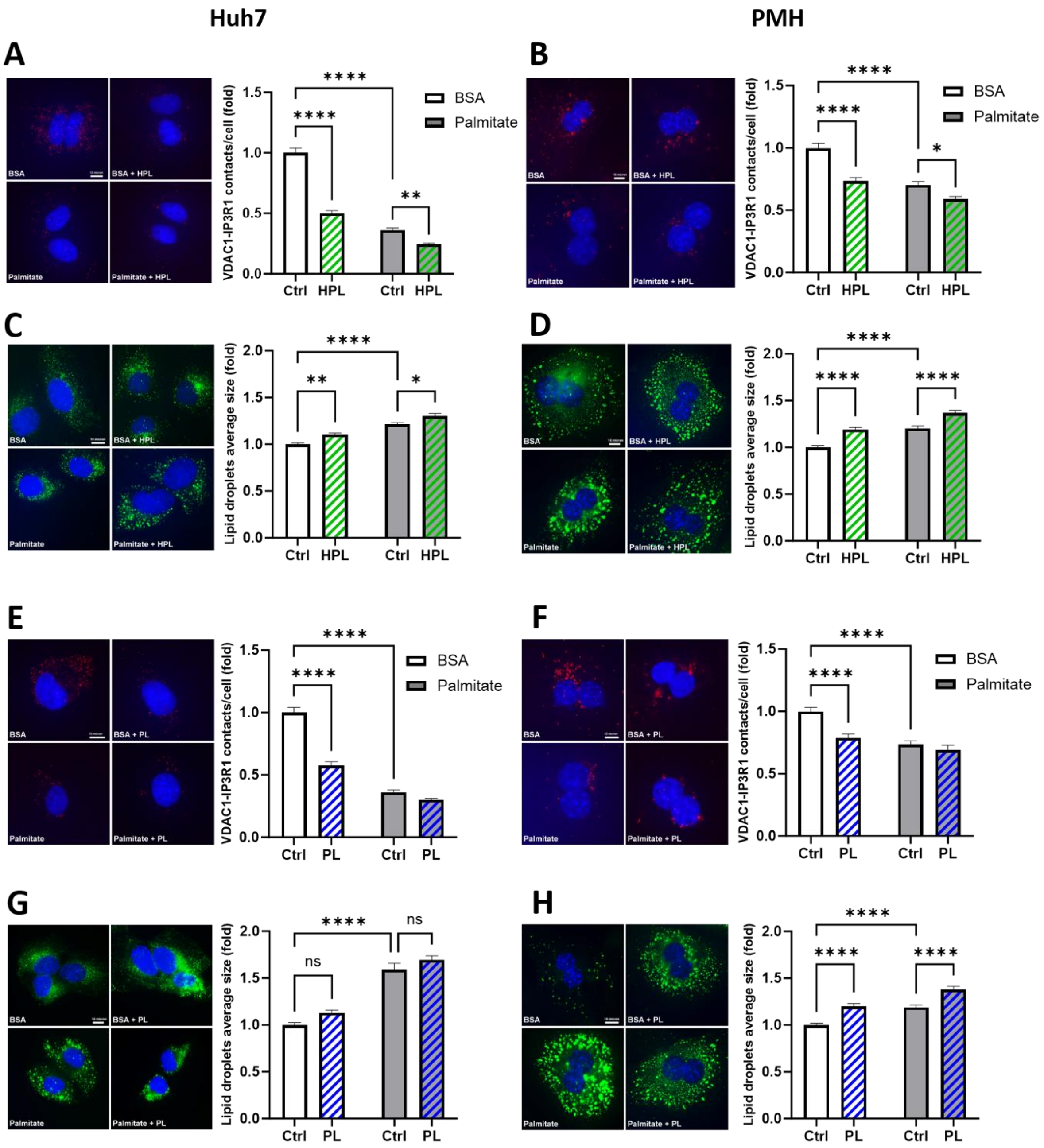


**Figure S4: HPL and PL induce lipid accumulation and disrupt ER-mitochondria interactions in hepatocytes.**

(A-H) Huh7 cells (A, C, E, G) or primary mouse hepatocytes (PMH, B, D, F, H) were treated for 16h with metabolites (HPL or PL, green and blue dashed bar, respectively) or vehicles (empty bar), in basal situation (BSA, white) or with 100µM palmitate (grey). Results are expressed as a fold of control. Representative images (left, scale bar: 10 µm) and quantitative analysis (right) of IP3R1-VDAC1 dots/cell measured by *in situ* PLA in both Huh7 cells (A, E, N=3-4 experiments, n=50-66 cells) and PMH (B, F, N=3-4 experiments, n=80 cells). Representative images (left, scale bar: 10 µm) and quantitative analysis (right) of lipid droplets average size in both Huh7 cells (C, G, N=3-4 experiments, n=36-90 cells) and PMH (D, H, N=3-4 experiments, n=80 cells) measured by BODIPY staining. The effects of treatments on lipid droplets number were illustrated in Figure S1E-S1H. Statistical analyses: Two Way Anova followed by Tukey's multiple comparison test: *p <0.05; **p<0.01; ***p <0.001; ****p <0.0001.
