## Supplemental Table S1_S2 for "N-acetyl-phenylalanine induces hepatic steatosis in MASLD by disrupting ER-mitochondria calcium coupling and mitochondrial lipid oxidation"

**Supplemental Table S1: Metabolite derived from phenylalanine and tyrosine metabolism identified using untargeted metabolomic analysis in 156 well-characterized participants with hepatic fat content assessment using magnetic resonance imaging (MRI) proton density fat fraction (PDFF).** Green indicates significant difference (p≤0.05) between the groups shown; GREEN indicates a ratio < 1. Red indicates significant difference (p≤0.05) between the groups shown; RED indicates a ratio > 1. Pink indicates significant fold change and significant correlation and median p-value significant.

| | | | | | | | | | | | | | % Filled values | | | | | |
|---|---|---|---|---|---|---|---|---|---|---|---|---|---|---|---|---|---|---|
| **Biochemical Name** | **Platform** | **Comp ID** | **KEGG** | **HMDB** | **PUBCHEM** | **Fold of Change NAFDL/non NAFLD** | **p-value** | **q-value** | **MRI PDFF AV** | **p-value** | **q-value** | **CORRELATION** | **NON** | **NAFLD** | **Non** | **NAFLD** | **Super Path way** | **Sub Pathway** |
| N-acetylphenylalanine | LC/MS neg | 33950 | C03519 | HMDB00512 | 74839 | 1,54 | 4,48E-07 | 1,92E-05 | | 1,18E-11 | 8,61E-10 | 0,5117 | 1,0105 | 1,555 | 100 | 100 | Amino Acid | Phenylalanine and Tyrosine Metabolism |
| 4-hydroxyphenylpyruvate | LC/MS neg | 1669 | C01179 | HMDB00707 | 979 | 1,34 | 8,04E-06 | 0,0001 | | 4,65E-07 | 5,87E-06 | 0,3929 | 0,989 | 1,3288 | 100 | 100 | Amino Acid | Phenylalanine and Tyrosine Metabolism |
| tyrosine | LC/MS pos | 1299 | C00082 | HMDB00158 | 6057 | 1,14 | 0,0005 | 0,0026 | | 9,99E-06 | 0,0001 | 0,3476 | 0,9905 | 1,1303 | 100 | 100 | Amino Acid | Phenylalanine and Tyrosine Metabolism |
| 3-(4-hydroxyphenyl)lactate | LC/MS neg | 32197 | C03672 | HMDB00755 | 9378 | 1,42 | 1,01E-06 | 2,74E-05 | | 2,24E-05 | 0,0001 | 0,3345 | 1,0002 | 1,4187 | 100 | 100 | Amino Acid | Phenylalanine and Tyrosine Metabolism |
| phenylpyruvate | LC/MS neg | 566 | C00166 | HMDB00205 | 997 | 1,34 | 0,0008 | 0,0036 | | 0,0001 | 0,0005 | 0,3078 | 0,9848 | 1,3179 | 100 | 100 | Amino Acid | Phenylalanine and Tyrosine Metabolism |
| phenol sulfate | LC/MS neg | 32553 | C02180 | HMDB60015 | 74426 | 1,47 | 0,0027 | 0,0084 | | 0,0005 | 0,0015 | 0,2793 | 1,2015 | 1,7715 | 100 | 100 | Amino Acid | Phenylalanine and Tyrosine Metabolism |
| 2-hydroxyphenylacetate | LC/MS polar | 1432 | C05852 | HMDB00669 | 11970 | 1,3 | 0,0027 | 0,0084 | | 0,0005 | 0,0018 | 0,2756 | 1,024 | 1,3311 | 97 | 100 | Amino Acid | Phenylalanine and Tyrosine Metabolism |
| N-acetyltyrosine | LC/MS neg | 32390 | | HMDB00866 | 68310 | 0,51 | 0,0007 | 0,0032 | | 0,0012 | 0,0034 | 0,258 | 2,9517 | 1,4979 | 100 | 100 | Amino Acid | Phenylalanine and Tyrosine Metabolism |
| 4-hydroxyphenylacetate | GC/MS | 541 | C00642 | HMDB00020 | 127 | 1,61 | 0,0176 | 0,0365 | | 0,0014 | 0,0037 | 0,256 | 0,9815 | 1,5805 | 72 | 81 | Amino Acid | Phenylalanine and Tyrosine Metabolism |
| phenylalanine | LC/MS pos | 64 | C00079 | HMDB00159 | 6140 | 1,1 | 0,0019 | 0,0064 | | 0,0025 | 0,0062 | 0,2417 | 0,9969 | 1,1002 | 100 | 100 | Amino Acid | Phenylalanine and Tyrosine Metabolism |
| phenyllactate (PLA) | LC/MS neg | 22130 | C05607 | HMDB00779 | 3848 | 1,2 | 0,0005 | 0,0024 | | 0,0096 | 0,0172 | 0,2081 | 1,0573 | 1,2708 | 100 | 100 | Amino Acid | Phenylalanine and Tyrosine Metabolism |
| 3-methoxytyramine sulfate | LC/MS neg | 44618 | | | | 1,21 | 0,1273 | 0,1544 | | 0,0127 | 0,0215 | 0,2005 | 1,099 | 1,3284 | 99 | 100 | Amino Acid | Phenylalanine and Tyrosine Metabolism |
| vanillylmandelate (VMA) | LC/MS neg | 1567 | C05584 | HMDB00291 | 1245 | 1,14 | 0,0165 | 0,0347 | | 0,0452 | 0,0588 | 0,1616 | 1,0362 | 1,1812 | 100 | 100 | Amino Acid | Phenylalanine and Tyrosine Metabolism |
| o-cresol sulfate | LC/MS neg | 36845 | | | 11615528 | 1,77 | 0,0945 | 0,1242 | | 0,0989 | 0,1093 | 0,1335 | 1,2305 | 2,1734 | 100 | 100 | Amino Acid | Phenylalanine and Tyrosine Metabolism |
| 3-[3-(sulfooxy)phenyl]propanoic acid | LC/MS neg | 45415 | | | 187488 | 1,42 | 0,547 | 0,3928 | | 0,3555 | 0,2714 | 0,075 | 1,175 | 1,6636 | 69 | 69 | Amino Acid | Phenylalanine and Tyrosine Metabolism |
| gentisate | LC/MS neg | 18280 | C00628 | HMDB00152 | 3469 | 0,96 | 0,7738 | 0,4752 | | 0,5034 | 0,3369 | 0,0543 | 1,211 | 1,1595 | 96 | 97 | Amino Acid | Phenylalanine and Tyrosine Metabolism |

| | | | | | | | | | | | | | | | | | | |
|---|---|---|---|---|---|---|---|---|---|---|---|---|---|---|---|---|---|---|
| phenylacetylglutamine | LC/MS pos | 35126 | C04148 | HMDB06344 | 92258 | 1,02 | 0,8549 | 0,5033 | | 0,5505 | 0,357 | 0,0485 | 1,097 | 1,1138 | 100 | 100 | Amino Acid | Phenylalanine and Tyrosine Metabolism |
| 3-(3-hydroxyphenyl)propionate | LC/MS neg | 35635 | C11457 | HMDB00375 | 91 | 1,17 | 0,8579 | 0,5033 | | 0,7943 | 0,4392 | 0,0212 | 1,6259 | 1,9 | 92 | 89 | Amino Acid | Phenylalanine and Tyrosine Metabolism |
| p-cresol sulfate | LC/MS neg | 36103 | C01468 | HMDB11635 | 4615423 | 0,8 | 0,3953 | 0,3187 | | 0,9528 | 0,4843 | -0,0048 | 1,1929 | 0,9524 | 100 | 100 | Amino Acid | Phenylalanine and Tyrosine Metabolism |
| 3-(4-hydroxyphenyl)propionate | LC/MS neg | 39587 | C01744 | HMDB02199 | 10394 | 1,19 | 0,4822 | 0,3623 | | 0,9034 | 0,4745 | -0,0099 | 1,0026 | 1,1933 | 60 | 69 | Amino Acid | Phenylalanine and Tyrosine Metabolism |
| 3-methoxytyrosine | LC/MS pos | 12017 | | HMDB01434 | 1670 | 1,05 | 0,6944 | 0,4537 | | 0,7478 | 0,4223 | -0,0261 | 1,0078 | 1,0576 | 100 | 100 | Amino Acid | Phenylalanine and Tyrosine Metabolism |
| Thyroxine | LC/MS pos | 46079 | C01829 | HMDB01918 | 5819 | 0,88 | 0,4943 | 0,3687 | | 0,2345 | 0,2039 | -0,0964 | 0,9259 | 0,818 | 76 | 72 | Amino Acid | Phenylalanine and Tyrosine Metabolism |
| 3-phenylpropionate (hydrocinnamate) | LC/MS neg | 15749 | C05629 | HMDB00764 | 107 | **0,64** | 0,0292 | 0,0525 | | 0,1233 | 0,126 | -0,1247 | 1,3395 | 0,8612 | 100 | 100 | Amino Acid | Phenylalanine and Tyrosine Metabolism |
| phenylacetate | LC/MS neg | 15958 | C07086 | HMDB00209 | 999 | 0,92 | 0,4273 | 0,336 | | 0,0485 | 0,0626 | -0,1592 | 0,9113 | 0,8377 | 72 | 64 | Amino Acid | Phenylalanine and Tyrosine Metabolism |

**Supplemental Table S2: Metabolite derived from phenylalanine and tyrosine metabolism identified using untargeted metabolomic analysis in 156 well-characterized participants with hepatic fat content assessment using biopsies.** Non-colored text and cell indicate that mean values are not significantly different for that comparison. Blue cell indicates significant difference (p≤0.05). Light Blue cell indicates p>0.05, p<0.10.

| | | | | | | | Correlations | Steatosis | | | |
|---|---|---|---|---|---|---|---|---|---|---|---|
| Sub Pathway | Biochemical Name | Platform | Comp ID | KEGG | HMDB | PUBCHEM | Steatosis | p-value | q-value | CORRELATION | Super Pathway |
| Phenylalanine and Tyrosine Metabolism | N-acetylphenylalanine | LC/MS neg | 33950 | C03519 | HMDB00512 | 74839 | | 0,008 | 0,2233 | 0,2116 | Amino Acid |
| | phenyllactate (PLA) | LC/MS neg | 22130 | C05607 | HMDB00779 | 3848 | | 0,0307 | 0,3648 | 0,173 | Amino Acid |
| | N-acetyltyrosine | LC/MS neg | 32390 | | HMDB00866 | 68310 | | 0,0131 | 0,2641 | 0,1982 | Amino Acid |
| | 3-(4-hydroxyphenyl)lactate | LC/MS neg | 32197 | C03672 | HMDB00755 | 9378 | | 0,0318 | 0,3723 | 0,172 | Amino Acid |
| | 3-methoxytyramine sulfate | LC/MS neg | 44618 | | | | | 0,0151 | 0,2973 | 0,1942 | Amino Acid |
| | dopamine sulfate (2) | LC/MS neg | 48407 | | | | | 0,0247 | 0,3456 | 0,1798 | Amino Acid |
| | phenylalanine | LC/MS pos | 64 | C00079 | HMDB00159 | 6140 | | 0,4236 | 0,758 | 0,0645 | Amino Acid |
| | phenylpyruvate | LC/MS neg | 566 | C00166 | HMDB00205 | 997 | | 0,0514 | 0,4505 | 0,1562 | Amino Acid |
| | phenylacetate | LC/MS neg | 15958 | C07086 | HMDB00209 | 999 | | 0,6122 | 0,7882 | -0,0409 | Amino Acid |
| | 4-hydroxyphenylacetate | LC/MS neg | 541 | C00642 | HMDB00020 | 127 | | 0,7519 | 0,8018 | 0,0255 | Amino Acid |
| | phenylacetylglutamine | LC/MS pos | 35126 | C04148 | HMDB06344 | 92258 | | 0,2689 | 0,6969 | 0,0891 | Amino Acid |
| | tyrosine | LC/MS pos | 1299 | C00082 | HMDB00158 | 6057 | | 0,1599 | 0,617 | 0,1131 | Amino Acid |
| | 4-hydroxyphenylpyruvate | LC/MS neg | 1669 | C01179 | HMDB00707 | 979 | | 0,6085 | 0,7882 | -0,0413 | Amino Acid |
| | phenol sulfate | LC/MS neg | 32553 | C02180 | HMDB60015 | 74426 | | 0,5893 | 0,7882 | 0,0436 | Amino Acid |
| | p-cresol sulfate | LC/MS neg | 36103 | C01468 | HMDB11635 | 4615423 | | 0,837 | 0,8209 | 0,0166 | Amino Acid |
| | o-cresol sulfate | LC/MS neg | 36845 | | | 11615528 | | 0,1008 | 0,5454 | 0,1319 | Amino Acid |
| | vanillylmandelate (VMA) | LC/MS neg | 1567 | C05584 | HMDB00291 | 1245 | | 0,6481 | 0,7916 | -0,0368 | Amino Acid |
| | 3-methoxytyrosine | LC/MS pos | 12017 | | HMDB01434 | 1670 | | 0,4456 | 0,7585 | -0,0615 | Amino Acid |
| | homovanillate (HVA) | LC/MS neg | 1101 | C05582 | HMDB00118 | 1738 | | 0,6996 | 0,7966 | -0,0311 | Amino Acid |
| | gentisate | LC/MS neg | 18280 | C00628 | HMDB00152 | 3469 | | 0,8614 | 0,8288 | 0,0141 | Amino Acid |
| | 3-(3-hydroxyphenyl)propionate | LC/MS neg | 35635 | C11457 | HMDB00375 | 91 | | 0,9956 | 0,8453 | -0,0004 | Amino Acid |
| | 3-(4-hydroxyphenyl)propionate | LC/MS neg | 39587 | C01744 | HMDB02199 | 10394 | | 0,9523 | 0,8356 | -0,0048 | Amino Acid |
| | 3-phenylpropionate (hydrocinnamate) | LC/MS neg | 15749 | C05629 | HMDB00764 | 107 | | 0,6853 | 0,7966 | 0,0327 | Amino Acid |
| | thyroxine | LC/MS pos | 46079 | C01829 | HMDB01918 | 5819 | | 0,4182 | 0,758 | 0,0653 | Amino Acid |
| | phenylacetylcarnitine | LC/MS pos | 48425 | | | | | 0,6043 | 0,7882 | -0,0418 | Amino Acid |
| | dopamine sulfate (1) | LC/MS neg | 48406 | | | | | 0,2453 | 0,6788 | 0,0936 | Amino Acid |
| | p-cresol-glucuronide* | LC/MS neg | 48841 | | HMDB11686 | 154035 | | 0,58 | 0,7882 | 0,0446 | Amino Acid |
| | tyramine O-sulfate | LC/MS neg | 48408 | | HMDB06409 | 153005 | | 0,9022 | 0,8347 | -0,0099 | Amino Acid |
| | vanillic alcohol sulfate | LC/MS neg | 48733 | | | | | 0,4068 | 0,7493 | -0,0669 | Amino Acid |
