## Supplemental Table S3_S4_S5 for "N-acetyl-phenylalanine induces hepatic steatosis in MASLD by disrupting ER-mitochondria calcium coupling and mitochondrial lipid oxidation"

Bacterial species significantly correlated with plasma NAPA level.
green: positive correlation, blue : negative correlation

| Bacterial Species | rho Correlation Coefficient | p,value | statistic | n | gp | Method |
|---|---|---|---|---|---|---|
| Actinomyces_sp._ICM39 | 0,248 | 0,043 | 2,06 | 69 | 2 | spearman |
| Anaerostipes_sp._CAG.276 | -0,270 | 0,027 | -2,26 | 69 | 2 | spearman |
| Bifidobacterium_longum | -0,258 | 0,035 | -2,15 | 69 | 2 | spearman |
| Bifidobacterium_moukalabense | -0,259 | 0,035 | -2,16 | 69 | 2 | spearman |
| Catenibacterium_mitsuokai | 0,295 | 0,015 | 2,49 | 69 | 2 | spearman |
| Clostridium_acidurici | -0,256 | 0,037 | -2,13 | 69 | 2 | spearman |
| Clostridium_colicanis | -0,243 | 0,047 | -2,02 | 69 | 2 | spearman |
| Clostridium_sp._CAG.149 | -0,259 | 0,034 | -2,16 | 69 | 2 | spearman |
| Clostridium_sp._CAG.167 | 0,281 | 0,021 | 2,36 | 69 | 2 | spearman |
| Clostridium_sp._CAG.230 | -0,241 | 0,049 | -2,01 | 69 | 2 | spearman |
| Clostridium_sp._CAG.277 | -0,268 | 0,028 | -2,24 | 69 | 2 | spearman |
| Clostridium_sp._CAG.575 | -0,289 | 0,018 | -2,43 | 69 | 2 | spearman |
| Clostridium_sp._CAG.729 | -0,243 | 0,047 | -2,02 | 69 | 2 | spearman |
| Clostridium_sp._CAG.81 | -0,244 | 0,046 | -2,03 | 69 | 2 | spearman |
| Dialister_micraerophilus | -0,289 | 0,018 | -2,44 | 69 | 2 | spearman |
| Eggerthella_sp._CAG.1427 | -0,254 | 0,038 | -2,12 | 69 | 2 | spearman |
| Enterobacter_sp._MGH_33 | 0,244 | 0,047 | 2,03 | 69 | 2 | spearman |
| Enterococcus_faecalis | -0,271 | 0,027 | -2,27 | 69 | 2 | spearman |
| Eubacterium_brachy | 0,260 | 0,033 | 2,17 | 69 | 2 | spearman |
| Eubacterium_saphenum | -0,250 | 0,041 | -2,08 | 69 | 2 | spearman |
| Finegoldia_magna | -0,272 | 0,026 | -2,28 | 69 | 2 | spearman |
| Firmicutes_bacterium_CAG.270 | -0,348 | 0,004 | -2,99 | 69 | 2 | spearman |
| Firmicutes_bacterium_CAG.466 | -0,299 | 0,014 | -2,53 | 69 | 2 | spearman |
| Fusobacterium_periodonticum | -0,250 | 0,042 | -2,08 | 69 | 2 | spearman |
| Geitlerinema_sp._PCC_7407 | 0,323 | 0,008 | 2,75 | 69 | 2 | spearman |
| Lachnospiraceae_bacterium_oral_taxon_082 | 0,277 | 0,023 | 2,32 | 69 | 2 | spearman |
| Lactobacillus_farraginis | 0,246 | 0,044 | 2,05 | 69 | 2 | spearman |
| Odoribacter_laneus | 0,250 | 0,041 | 2,08 | 69 | 2 | spearman |
| Parabacteroides_johnsonii | -0,251 | 0,041 | -2,09 | 69 | 2 | spearman |
| Prevotella_bergensis | 0,315 | 0,009 | 2,68 | 69 | 2 | spearman |
| Prevotella_copri_CAG.164 | 0,282 | 0,021 | 2,37 | 69 | 2 | spearman |
| Prevotella_disiens | 0,245 | 0,045 | 2,04 | 69 | 2 | spearman |
| Prevotella_oris | 0,245 | 0,045 | 2,04 | 69 | 2 | spearman |
| Prevotella_saccharolytica | 0,246 | 0,045 | 2,05 | 69 | 2 | spearman |
| Prevotella_sp._ICM33 | 0,246 | 0,045 | 2,05 | 69 | 2 | spearman |
| Prevotella_sp._oral_taxon_299 | 0,245 | 0,045 | 2,04 | 69 | 2 | spearman |
| Prevotella_sp._oral_taxon_317 | 0,245 | 0,045 | 2,04 | 69 | 2 | spearman |
| Prevotella_timonensis | 0,245 | 0,045 | 2,04 | 69 | 2 | spearman |
| Ruegeria_sp._TM1040 | -0,243 | 0,047 | -2,02 | 69 | 2 | spearman |
| Ruminococcus_champanellensis | -0,244 | 0,046 | -2,03 | 69 | 2 | spearman |
| Tannerella_sp._CAG.51 | 0,280 | 0,022 | 2,35 | 69 | 2 | spearman |
| Variovorax_sp._CF313 | 0,287 | 0,019 | 2,41 | 69 | 2 | spearman |

**SUPPLEMENTAL TABLE S4**

Bacterial species significantly correlated with hepatic fat content measured by MRI-PDFF.
Green: positive correlation, blue: negative correlation

| Bacterial Species | rho Correlation Coefficient | p,value | statistic | n | gp | Method |
|---|---|---|---|---|---|---|
| Acholeplasma_palmae | -0,365 | 0,002 | -3,158 | 69 | 2 | spearman |
| Acinetobacter_sp._ATCC_27244 | -0,308 | 0,011 | -2,607 | 69 | 2 | spearman |
| Actinotalea_ferrariae | -0,284 | 0,020 | -2,390 | 69 | 2 | spearman |
| Alishewanella_jeotgali | -0,355 | 0,003 | -3,058 | 69 | 2 | spearman |
| Alistipes_finegoldii | -0,278 | 0,023 | -2,333 | 69 | 2 | spearman |
| Alistipes_putredinis_CAG.67 | -0,294 | 0,016 | -2,476 | 69 | 2 | spearman |
| Alistipes_sp._CAG.157 | -0,252 | 0,040 | -2,096 | 69 | 2 | spearman |
| Alistipes_sp._CAG.435 | 0,251 | 0,040 | 2,095 | 69 | 2 | spearman |
| Alistipes_sp._CAG.514 | -0,317 | 0,009 | -2,699 | 69 | 2 | spearman |
| Alistipes_sp._CAG.831 | -0,355 | 0,003 | -3,058 | 69 | 2 | spearman |
| Anaerococcus_lactolyticus | 0,299 | 0,014 | 2,522 | 69 | 2 | spearman |
| Anaerotruncus_colihominis | -0,411 | 0,001 | -3,634 | 69 | 2 | spearman |
| Anaerotruncus_sp._CAG.528 | -0,266 | 0,030 | -2,224 | 69 | 2 | spearman |
| Bacteroidales_bacterium_CF | -0,293 | 0,016 | -2,474 | 69 | 2 | spearman |
| Bacteroides_eggerthii | -0,339 | 0,005 | -2,906 | 69 | 2 | spearman |
| Bacteroides_helcogenes | -0,260 | 0,033 | -2,173 | 69 | 2 | spearman |
| Bacteroides_massiliensis | -0,308 | 0,011 | -2,608 | 69 | 2 | spearman |
| Bacteroides_nordii | -0,325 | 0,007 | -2,771 | 69 | 2 | spearman |
| Bacteroides_sp._CAG.443 | 0,310 | 0,011 | 2,627 | 69 | 2 | spearman |
| Bacteroides_sp._CAG.530 | 0,271 | 0,026 | 2,272 | 69 | 2 | spearman |
| Bacteroides_sp._CAG.702 | 0,302 | 0,013 | 2,550 | 69 | 2 | spearman |
| Bacteroides_sp._D22 | -0,288 | 0,018 | -2,421 | 69 | 2 | spearman |
| Bifidobacterium_longum_CAG.69 | 0,243 | 0,047 | 2,024 | 69 | 2 | spearman |
| Butyrivibrio_crossotus | -0,261 | 0,033 | -2,176 | 69 | 2 | spearman |
| Butyrivibrio_proteoclasticus | -0,260 | 0,034 | -2,170 | 69 | 2 | spearman |
| Catenibacterium_mitsuokai | 0,322 | 0,008 | 2,746 | 69 | 2 | spearman |
| Clostridiales_bacterium_BV3C26 | -0,313 | 0,010 | -2,661 | 69 | 2 | spearman |
| Clostridium_bolteae | 0,255 | 0,037 | 2,128 | 69 | 2 | spearman |
| Clostridium_sp._BNL1100 | -0,264 | 0,031 | -2,205 | 69 | 2 | spearman |
| Clostridium_sp._CAG.1013 | -0,337 | 0,005 | -2,882 | 69 | 2 | spearman |
| Clostridium_sp._CAG.1024 | -0,304 | 0,012 | -2,572 | 69 | 2 | spearman |
| Clostridium_sp._CAG.1193 | -0,311 | 0,010 | -2,636 | 69 | 2 | spearman |
| Clostridium_sp._CAG.127 | -0,365 | 0,002 | -3,160 | 69 | 2 | spearman |
| Clostridium_sp._CAG.169 | -0,282 | 0,021 | -2,373 | 69 | 2 | spearman |
| Clostridium_sp._CAG.221 | -0,288 | 0,018 | -2,425 | 69 | 2 | spearman |
| Clostridium_sp._CAG.230 | -0,301 | 0,013 | -2,543 | 69 | 2 | spearman |
| Clostridium_sp._CAG.242 | -0,378 | 0,002 | -3,294 | 69 | 2 | spearman |
| Clostridium_sp._CAG.265 | -0,313 | 0,010 | -2,661 | 69 | 2 | spearman |
| Clostridium_sp._CAG.269 | -0,305 | 0,012 | -2,584 | 69 | 2 | spearman |
| Clostridium_sp._CAG.277 | -0,269 | 0,028 | -2,248 | 69 | 2 | spearman |
| Clostridium_sp._CAG.299 | -0,382 | 0,001 | -3,335 | 69 | 2 | spearman |
| Clostridium_sp._CAG.302 | -0,293 | 0,016 | -2,469 | 69 | 2 | spearman |
| Clostridium_sp._CAG.307 | -0,275 | 0,024 | -2,307 | 69 | 2 | spearman |
| Clostridium_sp._CAG.349 | -0,334 | 0,006 | -2,855 | 69 | 2 | spearman |
| Clostridium_sp._CAG.352 | -0,358 | 0,003 | -3,093 | 69 | 2 | spearman |

| | | | | | | |
|---|---|---|---|---|---|---|
| **Clostridium_sp._CAG.354** | -0,306 | 0,012 | -2,591 | 69 | 2 | spearman |
| **Clostridium_sp._CAG.389** | -0,251 | 0,041 | -2,088 | 69 | 2 | spearman |
| **Clostridium_sp._CAG.411** | -0,251 | 0,041 | -2,090 | 69 | 2 | spearman |
| **Clostridium_sp._CAG.413** | -0,421 | 0,000 | -3,741 | 69 | 2 | spearman |
| **Clostridium_sp._CAG.43** | -0,292 | 0,016 | -2,465 | 69 | 2 | spearman |
| **Clostridium_sp._CAG.433** | -0,282 | 0,021 | -2,369 | 69 | 2 | spearman |
| **Clostridium_sp._CAG.448** | -0,331 | 0,006 | -2,831 | 69 | 2 | spearman |
| **Clostridium_sp._CAG.451** | -0,243 | 0,048 | -2,019 | 69 | 2 | spearman |
| **Clostridium_sp._CAG.452** | -0,271 | 0,027 | -2,270 | 69 | 2 | spearman |
| **Clostridium_sp._CAG.470** | -0,334 | 0,006 | -2,858 | 69 | 2 | spearman |
| **Clostridium_sp._CAG.505** | -0,309 | 0,011 | -2,622 | 69 | 2 | spearman |
| **Clostridium_sp._CAG.510** | -0,253 | 0,039 | -2,106 | 69 | 2 | spearman |
| **Clostridium_sp._CAG.557** | -0,274 | 0,025 | -2,293 | 69 | 2 | spearman |
| **Clostridium_sp._CAG.568** | -0,259 | 0,035 | -2,158 | 69 | 2 | spearman |
| **Clostridium_sp._CAG.594** | -0,337 | 0,005 | -2,882 | 69 | 2 | spearman |
| **Clostridium_sp._CAG.62** | -0,339 | 0,005 | -2,902 | 69 | 2 | spearman |
| **Clostridium_sp._CAG.7** | -0,373 | 0,002 | -3,240 | 69 | 2 | spearman |
| **Clostridium_sp._CAG.729** | -0,272 | 0,026 | -2,278 | 69 | 2 | spearman |
| **Clostridium_sp._CAG.75** | -0,332 | 0,006 | -2,840 | 69 | 2 | spearman |
| **Clostridium_sp._CAG.768** | -0,257 | 0,035 | -2,148 | 69 | 2 | spearman |
| **Clostridium_sp._CAG.780** | -0,324 | 0,007 | -2,765 | 69 | 2 | spearman |
| **Clostridium_sp._CAG.793** | -0,341 | 0,005 | -2,926 | 69 | 2 | spearman |
| **Clostridium_sp._CAG.81** | -0,244 | 0,046 | -2,030 | 69 | 2 | spearman |
| **Clostridium_sp._CAG.91** | -0,376 | 0,002 | -3,272 | 69 | 2 | spearman |
| **Clostridium_sp._CAG.914** | -0,361 | 0,003 | -3,120 | 69 | 2 | spearman |
| **Clostridium_sp._HGF2** | -0,246 | 0,045 | -2,042 | 69 | 2 | spearman |
| **Coprobacillus_sp._CAG.235** | -0,330 | 0,006 | -2,820 | 69 | 2 | spearman |
| **Enterobacter_sp._MGH_16** | 0,258 | 0,035 | 2,157 | 69 | 2 | spearman |
| **Erysipelotrichaceae_bacterium_5_2_54FAA** | -0,250 | 0,041 | -2,080 | 69 | 2 | spearman |
| **Eubacterium_saphenum** | -0,494 | 0,000 | -4,575 | 69 | 2 | spearman |
| **Eubacterium_siraeum** | -0,520 | 0,000 | -4,911 | 69 | 2 | spearman |
| **Eubacterium_sp._CAG.146** | -0,285 | 0,019 | -2,397 | 69 | 2 | spearman |
| **Eubacterium_sp._CAG.786** | -0,288 | 0,018 | -2,421 | 69 | 2 | spearman |
| **Firmicutes_bacterium_CAG.103** | -0,289 | 0,018 | -2,438 | 69 | 2 | spearman |
| **Firmicutes_bacterium_CAG.110** | -0,275 | 0,024 | -2,310 | 69 | 2 | spearman |
| **Firmicutes_bacterium_CAG.114** | -0,288 | 0,018 | -2,427 | 69 | 2 | spearman |
| **Firmicutes_bacterium_CAG.129** | -0,244 | 0,047 | -2,026 | 69 | 2 | spearman |
| **Firmicutes_bacterium_CAG.137** | -0,266 | 0,030 | -2,225 | 69 | 2 | spearman |
| **Firmicutes_bacterium_CAG.145** | -0,318 | 0,009 | -2,703 | 69 | 2 | spearman |
| **Firmicutes_bacterium_CAG.170** | -0,245 | 0,045 | -2,041 | 69 | 2 | spearman |
| **Firmicutes_bacterium_CAG.176** | -0,249 | 0,042 | -2,075 | 69 | 2 | spearman |
| **Firmicutes_bacterium_CAG.227** | -0,263 | 0,031 | -2,202 | 69 | 2 | spearman |
| **Firmicutes_bacterium_CAG.24** | -0,375 | 0,002 | -3,256 | 69 | 2 | spearman |
| **Firmicutes_bacterium_CAG.240** | -0,282 | 0,021 | -2,373 | 69 | 2 | spearman |
| **Firmicutes_bacterium_CAG.270** | -0,340 | 0,005 | -2,913 | 69 | 2 | spearman |
| **Firmicutes_bacterium_CAG.308** | -0,356 | 0,003 | -3,072 | 69 | 2 | spearman |
| **Firmicutes_bacterium_CAG.313** | -0,274 | 0,025 | -2,298 | 69 | 2 | spearman |
| **Firmicutes_bacterium_CAG.449** | -0,312 | 0,010 | -2,651 | 69 | 2 | spearman |
| **Firmicutes_bacterium_CAG.466** | -0,297 | 0,015 | -2,512 | 69 | 2 | spearman |
| **Firmicutes_bacterium_CAG.791** | -0,293 | 0,016 | -2,472 | 69 | 2 | spearman |
| **Firmicutes_bacterium_CAG.822** | -0,274 | 0,025 | -2,299 | 69 | 2 | spearman |
| **Firmicutes_bacterium_CAG.882** | -0,289 | 0,018 | -2,432 | 69 | 2 | spearman |
| **Firmicutes_bacterium_CAG.884** | -0,278 | 0,023 | -2,337 | 69 | 2 | spearman |

| | | | | | | |
|---|---|---|---|---|---|---|
| Labrenzia_alexandrii | 0,267 | 0,029 | 2,234 | 69 | 2 | spearman |
| Lachnospiraceae_bacterium_2_1_46FAA | 0,258 | 0,035 | 2,151 | 69 | 2 | spearman |
| Lachnospiraceae_bacterium_5_1_63FAA | 0,271 | 0,026 | 2,273 | 69 | 2 | spearman |
| Lachnospiraceae_bacterium_CAG.25 | 0,261 | 0,033 | 2,182 | 69 | 2 | spearman |
| Lachnospiraceae_bacterium_oral_taxon_082 | 0,313 | 0,010 | 2,659 | 69 | 2 | spearman |
| Mycoplasma_putrefaciens | -0,327 | 0,007 | -2,786 | 69 | 2 | spearman |
| Mycoplasma_sp._CAG.472 | -0,279 | 0,022 | -2,347 | 69 | 2 | spearman |
| Mycoplasma_sp._CAG.776 | -0,316 | 0,009 | -2,685 | 69 | 2 | spearman |
| Oscillibacter_sp._CAG.155 | -0,251 | 0,040 | -2,092 | 69 | 2 | spearman |
| Owenweeksia_hongkongensis | -0,282 | 0,021 | -2,368 | 69 | 2 | spearman |
| Parabacteroides_distasonis | -0,315 | 0,010 | -2,672 | 69 | 2 | spearman |
| Parabacteroides_sp._CAG.2 | 0,284 | 0,020 | 2,391 | 69 | 2 | spearman |
| Phaeospirillum_fulvum | -0,259 | 0,034 | -2,165 | 69 | 2 | spearman |
| Prevotella_bergensis | 0,250 | 0,041 | 2,085 | 69 | 2 | spearman |
| Prevotella_bivia | 0,324 | 0,008 | 2,759 | 69 | 2 | spearman |
| Prevotella_bryantii | 0,324 | 0,007 | 2,765 | 69 | 2 | spearman |
| Prevotella_disiens | 0,249 | 0,042 | 2,076 | 69 | 2 | spearman |
| Prevotella_intermedia | 0,254 | 0,038 | 2,119 | 69 | 2 | spearman |
| Prevotella_maculosa | 0,268 | 0,028 | 2,241 | 69 | 2 | spearman |
| Prevotella_marshii | 0,331 | 0,006 | 2,826 | 69 | 2 | spearman |
| Prevotella_nigrescens | 0,323 | 0,008 | 2,748 | 69 | 2 | spearman |
| Prevotella_oris | 0,249 | 0,042 | 2,076 | 69 | 2 | spearman |
| Prevotella_pallens | 0,344 | 0,004 | 2,954 | 69 | 2 | spearman |
| Prevotella_saccharolytica | 0,250 | 0,041 | 2,085 | 69 | 2 | spearman |
| Prevotella_sp._BV3P1 | 0,291 | 0,017 | 2,452 | 69 | 2 | spearman |
| Prevotella_sp._CAG.1031 | 0,277 | 0,023 | 2,327 | 69 | 2 | spearman |
| Prevotella_sp._CAG.1092 | 0,364 | 0,002 | 3,149 | 69 | 2 | spearman |
| Prevotella_sp._CAG.255 | 0,273 | 0,025 | 2,289 | 69 | 2 | spearman |
| Prevotella_sp._ICM33 | 0,250 | 0,041 | 2,085 | 69 | 2 | spearman |
| Prevotella_sp._oral_taxon_299 | 0,249 | 0,042 | 2,076 | 69 | 2 | spearman |
| Prevotella_sp._oral_taxon_317 | 0,249 | 0,042 | 2,076 | 69 | 2 | spearman |
| Prevotella_timonensis | 0,249 | 0,042 | 2,076 | 69 | 2 | spearman |
| Pseudoalteromonas_undina | -0,333 | 0,006 | -2,851 | 69 | 2 | spearman |
| Pseudomonas_sp._VLB120 | 0,269 | 0,028 | 2,255 | 69 | 2 | spearman |
| richness | -0,342 | 0,005 | -2,931 | 69 | 2 | spearman |
| Roseburia_sp._CAG.197 | -0,254 | 0,038 | -2,119 | 69 | 2 | spearman |
| Roseburia_sp._CAG.303 | -0,267 | 0,029 | -2,230 | 69 | 2 | spearman |
| Ruminococcus_bromii | -0,362 | 0,003 | -3,130 | 69 | 2 | spearman |
| Ruminococcus_flavefaciens | 0,275 | 0,024 | 2,308 | 69 | 2 | spearman |
| Ruminococcus_sp._CAG.108 | -0,242 | 0,049 | -2,010 | 69 | 2 | spearman |
| Ruminococcus_sp._CAG.330 | -0,314 | 0,010 | -2,669 | 69 | 2 | spearman |
| Ruminococcus_sp._CAG.403 | -0,324 | 0,008 | -2,758 | 69 | 2 | spearman |
| Ruminococcus_sp._CAG.55 | -0,285 | 0,020 | -2,394 | 69 | 2 | spearman |
| Ruminococcus_sp._CAG.60 | -0,261 | 0,033 | -2,180 | 69 | 2 | spearman |
| Ruminococcus_sp._CAG.624 | -0,262 | 0,033 | -2,185 | 69 | 2 | spearman |
| Subdoligranulum_sp._4_3_54A2FAA | -0,356 | 0,003 | -3,075 | 69 | 2 | spearman |
| Syntrophus_aciditrophicus | 0,299 | 0,014 | 2,522 | 69 | 2 | spearman |
| Tannerella_sp._6_1_58FAA_CT1 | -0,295 | 0,015 | -2,489 | 69 | 2 | spearman |
| Tannerella_sp._CAG.51 | 0,320 | 0,008 | 2,724 | 69 | 2 | spearman |
| Veillonella_parvula | 0,281 | 0,021 | 2,360 | 69 | 2 | spearman |

**SUPPLEMENTAL TABLE S5**

| **Primers** | | | |
|---|---|---|---|
| ***Genes* - Species** | **Gene ID** | **Forward** | **Reverse** |
| *Acat1* - Mouse | 110446 | TGAAGCATTCAGTGTGGTTG | GCAAATACTAGCCAGACCGA |
| *Apob* - Mouse | 238055 | CCTCTCCTGGGTGTTCTAGAC | GTTTCTCCAGATCCTTGCAC |
| *Atgl* - Mouse | 66853 | TCATCCAGGCCAATCTCTGC | TGGATGTTGGTGGAGCTGTC |
| *Cd36* - Mouse | 12491 | GTCCTGGCTGTGTTTGGAGG | AAGATCCAAAACTGTCTGTA |
| *Chrebp* - Mouse | 58805 | AGCTCAACGCTGCCATCAAC | CTGAACACCCAGAACTTCCA |
| *Cidea* - Mouse | 12683 | CAGAGGAGTTCTTTCAGACC | GTATAGGTCGAAGGTGACTC |
| *Cideb* - Mouse | 12684 | CGTGTCTGTGATCATAAGCG | CGTATGACAACATCCCACTC |
| *Cpt1a* - Mouse | 12894 | ACGTATGAGGCTTCCATGAC | GGTGAGTCGACTGCCAGATA |
| *Dgat2* - Mouse | 67800 | TGGGTCCAGAAGAAGTTCCAGAAGTA | ACCTCAGTCTCTGGAAGGCCAAAT |
| *Hsl* - Mouse | 16835 | GTGTGTCAGTGCCTATTCAG | GTCAGCTTCTTCAAGGTATC |
| *Ldlr* - Mouse | 16890 | GTTGATTCCAAACTCCACTC | TTCACATCTGAACCCGTGAG |
| *Mttp* - Mouse | 17777 | GGAGAAGTAACCTGAACATC | ACAGGCCTAAGCTGAACATC |
| *Srebp1c* - Mouse | 20787 | ACGGAGCCATGGATTGCACA | AAGGGTGCAGGTGTCACCTT |
| *Tbp* - Mouse | 21374 | TGGTGTGCACAGGAGCCAAG | TTCACATCACAGCTCCCCAC |
| *SLC7A5* - Human | 8140 | CTTCAGCTTCTTCAACTGGC | AGGATGATGGTGAAGCCGAT |
| *TBP* - Human | 6908 | AGACCATTGCACTTCGTGCC | CCTGTGCACACCATTTTCCC |
